# Male-specific sNPF peptidergic circuits control energy balance for mating duration through neuron-glia interactions

**DOI:** 10.1101/2024.10.17.618859

**Authors:** Xiaoli Zhang, Hongyu Miao, Dayeon Kang, Dongyu Sun, Woo Jae Kim

## Abstract

This study reveals a sexually dimorphic sNPF-sNPF-R circuit in the *Drosophila* brain that regulates energy balance behavior, specifically the shorter-mating-duration (SMD) response to sexual experience. sNPF is predominantly expressed in neurons, while sNPF-R is expressed in both neurons and glial cells, particularly astrocyte-like glia (ALG) in a subset of cells outside the mushroom body (MB) termed "Rishi cells" (RS cells). Sexual experience induces global alterations in calcium signaling and synaptic plasticity within this circuit, with sNPF-R expressing neurons and glia in RS cells playing a critical role in encoding sexual experience-related information into energy balance behavior related to mating duration. Neuronal glucose metabolism, specifically the *Tret1l* transporter, is essential for maintaining the high calcium levels and proper function of sNPF neurons near RS cells. This novel and intricate neuron-glia sNPF-sNPF-R network acts as critical circuits within the *Drosophila* brain for processing interval timing behaviors, highlighting the dynamic interplay between metabolism, neural circuits, and behavior in regulating energy balance.

## INTRODUCTION

In the fruit fly *Drosophila melanogaster*, a sophisticated interplay of internal states and neural circuits regulates energy balance (Kim et al. 2024). These states, which include hunger and satiety signals, are integrated by specific groups of neurons that modulate feeding behavior and other energy-related activities. The internal states of energy balance in *Drosophila* are intricately linked to its physiology, behavior, and adaptation to environmental changes (Lee and Wu 2020; Münch et al. 2020; Chatterjee and Perrimon 2021; Reis 2023). This intricate system allows for a dynamic response to the fly’s nutritional needs, ensuring energy homeostasis is maintained.

For male *Drosophila*, as well as males across the animal kingdom, energy balance is crucial for maintaining reproductive health (Redman 2006; Gao and Horvath 2008; Morton et al. 2014; Gáliková et al. 2015; Koyama et al. 2021). This balance is achieved through a complex interplay of behavioral, physiological, and molecular mechanisms that ensure the organism has sufficient energy to support both daily activities and reproductive efforts. When energy intake is high, and expenditure is low, males can invest more in reproductive activities such as courtship, mating, and producing gametes. Conversely, during periods of energy scarcity, the investment in reproduction is often reduced to prioritize survival. For instance, when male *Drosophila* have prior mating experience, they tend to shorten the duration of copulation as a strategy to optimize energy allocation for reproductive activities (Kim et al. 2016; Wong et al. 2019; Lee et al. 2022; Lee et al. 2023; Kim et al. 2024; T. Zhang et al. 2024).

The complexity of the nervous system is not solely attributed to the intricate connections between neurons, but also to diverse glial cells that provide support and regulate the operation of neurons (Haydon 2001; Fields and Stevens-Graham 2002). The neuron-glia network is a complex biological system that evolved to effectively store and manipulate information for behavior (Laming et al. 2000). Neuron-glia interactions play a crucial role in the formation and maintenance of working memory (Bains and Oliet 2007; Vignoli et al. 2016; Pittà and Brunel 2022; Silva et al. 2022), the regulation of brain homeostasis and degeneration (Kettenmann et al. 1996), synaptic transmission (Haydon 2001; Perea and Araque 2010; Backer and Kadow 2022), the preservation of synaptic plasticity (Vernadakis 1996; Magistretti 2006; Stogsdill and Eroglu 2017), and the emergence and transition of internal brain states (Fields and Stevens-Graham 2002; Logan 2017; Verdugo et al. 2019).

Especially, the communication and interaction between networks of neurons and glial cells are essential for the maintenance of the brain’s internal states. Neuropeptides and their receptors have a crucial part in the complex communication that takes place within neuron-glia networks, making a considerable contribution to the maintenance and functioning of the central nervous system (CNS) both in human and fruit fly (Clasadonte and Prevot 2018; Nässel and Zandawala 2019; Nässel and Zandawala 2022). Neuropeptides play a crucial role in controlling synaptic plasticity, which is the ability of synapses to undergo lasting changes in their strength, either by increasing it or decreasing it. This process is necessary for learning and memory. One example is brain-derived neurotrophic factor (BDNF), which is a neuropeptide that enhances the ability of synapses to change and helps neurons survive (Sasi et al. 2017).

The short neuropeptide F (sNPF) and its receptor (sNPF-R) system in *Drosophila melanogaster* is a valuable model for studying the role and importance of neuropeptide signaling in human biology. sNPF is a neuropeptide that has a crucial function in controlling multiple physiological processes in *Drosophila*, including feeding behavior, aggression, sleep, and reproduction (Mertens et al. 2002; Lee et al. 2004; Lee et al. 2008; Nässel et al. 2008; Nässel and Wegener 2011; Chen et al. 2013; Knapek et al. 2013; Shang et al. 2013). The sNPF system exhibits notable resemblances to the neuropeptide Y (NPY) system in humans (Nässel and Wegener 2011). NPY is a neuropeptide that is extensively present in the human central nervous system and has a crucial function in controlling several physiological processes, such as feeding behavior, stress response, and energy metabolism (Nässel and Wegener 2011). The NPY system comprises NPY and its receptors (NPYRs), which play a role in transmitting the effects of NPY to target tissues (Medeiros and Turner 1996; Comeras et al. 2019).

Although there has been significant research on the *Drosophila* sNPF, the specific mechanisms by which sNPF-R controls metabolism through the neuropeptidergic system in the brain, as well as its interactions with neuron-glia networks, are not well understood. Here we show that male-specific sNPF to sNPF-R circuits in the brain play a crucial role in regulating energy balance behaviors that are affected by sexual experience. Our study demonstrates that there is a specific group of neuron-glia cells that express sNPF and sNPF-R which can integrate information from experiences and use it to modify the internal states of the brain.

## RESULTS

### sNPF-to-sNPF-R circuits are composed of functional neuron-glia network

In our previous work, we described the phenomenon of sexually experienced males reducing their investment in mating as a means to balance energy expenditure. This behavior has been termed shorter-mating-duration (SMD) (Kim et al. 2016; Lee et al. 2023). We utilize SMD as a behavioral platform to investigate the involvement of sNPF circuits in the neuromodulation of brain metabolism through the neuron-glia network, as sNPF is specifically associated with SMD behavior among neuropeptides expressed in 150 clock cells in the brain (Kim et al. 2013) (Fig. 1A-C) (Fig. S1A for genetic control).

**Figure 1.**
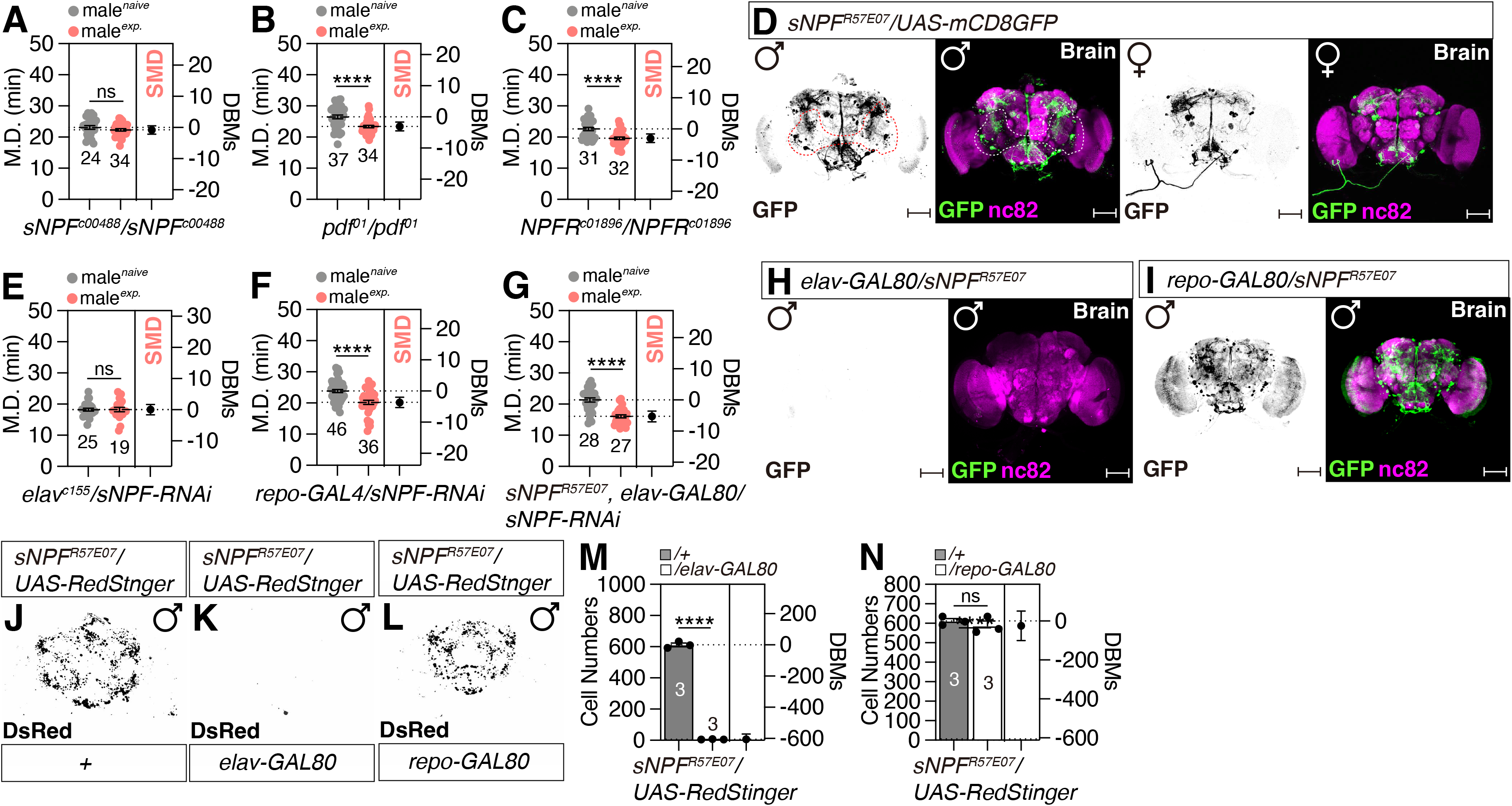
sNPF-to-sNPF-R circuits are composed of functional neuron-glia network. (A-C) SMD assays for *sNPF, pdf* and *NPFR* mutant. In the mating duration (MD) assays, light grey data points denote males that were sexually naïve, whereas pink data points signify males that were sexually experienced. The dot plots represent the MD of each male fly. The numerical values beneath the dot plots indicate the count of male flies that mated successfully. The mean value and standard error are labeled within the dot plot (black lines). M.D represent mating duration. DBMs represent the ’difference between means’ for the evaluation of estimation statistics (See **STAR** 11 **METHODS**). Asterisks represent significant differences, as revealed by the unpaired Student’s t test, and ns represents non-significant differences (*\*p<0.05, **p<0.01, ***p< 0.001, ****p< 0.0001*). Consequently, data points on graphs marked with asterisks indicate that SMD behavior remain unaltered or within normal parameters, whereas those labeled with ’ns’ signify that SMD behavior have been perturbed due to mutations or genetic alterations in the respective strains. For detailed methods, see the **STAR** 11 **METHODS** for a detailed description of the MD assays used in this study. In the framework of our investigation, the routine application of internal controls is employed for the vast majority of experimental procedures, as delineated in the "**Mating Duration Assay**" and "**Statistical Tests**" subsections of the **STAR** 11 **METHODS** section. The identical analytical approach employed for the MD assays is maintained for the subsequent data presented. (D) Brain of male and female flies expressing *sNPF^R57E07^*together with *UAS-CD8tdGFP* were immunostained with anti-GFP (green) and nc82 (magenta) antibodies. Scale bars represent 100 μm. For detailed methods, see the STAR 17 METHODS for a detailed description of the immunostaining procedure used in this study. (E-G) SMD assays of *elav^c155^* (E), *repo-GAL4* (F) and *elav-GAL80*, *sNPF^R57E07^* (G)-mediated knockdown of *sNPF via sNPF-RNAi.* (H-I) Brain of male flies expressing, *elav-GAL80, sNPF^R57E07^*(H) and *repo-GAL80, sNPF^R57E07^*(I) with *UAS-CD8tdGFP* were immunostained with anti-GFP (green) and nc82 (magenta) antibodies. (J-L) Brain of male flies expressing *sNPF^R57E07^* (J), *elav-GAL80*, *sNPF^R57E07^*(K), *repo-GAL80*, *sNPF^R57E07^* (L) with *UAS-Redstinger* were immunostained with anti-DsRed (gray). Scale bars represent 100 μm. (M-N) Quantification of cell number. The flies are described as (J-L). Bars represent the mean *sNPF^R57E07^* (gray column) and, *elav-GAL80*, *sNPF^R57E07^* (white column) cell number fluorescence level with error bars representing SEM (M). Bars represent the mean *sNPF^R57E07^* (gray column) and *repo-GAL80*, *sNPF^R57E07^* (white column) cell number fluorescence level with error bars representing SEM (N). Asterisks represent significant differences, as revealed by the Student’s *t* test and ns represents non-significant difference (*\*p<0.05, **p<0.01, ***p< 0.001, ****p< 0.0001*). See the **STAR** 11 **METHODS** for a detailed description of the particle analysis used in this study.

The *sNPF-GAL4* driver *sNPF^R57E07^* is chosen to replicate the natural expression patterns of sNPF, which are similar to those observed in the previously described *sNPF^2.5^-GAL4* (Lee et al. 2009) (Fig. 1D and Fig. S1B). The expression of the *sNPF^R57E07^* driver is more prominent in the male brain compared to the female brain (Fig. 1D and Fig. S1C). Knocking down sNPF using RNAi specifically in neurons, but not in glial cells, disrupts SMD behavior (Fig. 1E-F and for genetic control Fig. S1D). Additionally, expressing *sNPF-RNAi* only in cells labeled with *sNPF^R57E07^*, except for neurons, does not disrupt SMD (Fig. 1G).

Expression of GAL80 in neurons, but not in glial cells, can completely eliminate transgene expression in around 600 sNPF-positive cells identified by the *sNPF^R57E07^* driver (Fig. 1H-N). These data suggest that the presence of sNPF in the group of neurons is necessary and sufficient for producing SMD.

The *Drosophila* neuropeptide F receptor (sNPF-R) was first cloned over a decade ago (Mertens et al. 2002), yet detailed characterization of its expression pattern and circuit-level functions has remained an area of investigation. It has been observed that neuronal sNPF exerts a modulatory effect on energy homeostasis by activating sNPF-R in intestinal enterocytes, potentially contributing to the maintenance of gut epithelial integrity. This, in turn, may trigger a neural sensorimotor pathway that governs energy balance, potentially through the mediation of mechanisms related to disease tolerance (Shen et al. 2016).

The targeted knockdown of sNPF-R in both neuronal and glial cells resulted in the disruption of SMD behavior (Fig. 2A-B and Fig. S1E for genetic control). Out of the 14 GAL4 drivers that were available (Pfeiffer et al. 2008; Arbel et al. 2019), *sNPF-R^R64H09^* was selected as a putative sNPF-R driver due to its extensive labeling within the most richly innervated region of the central brain (CB) (Table S1). The expression of neuronal GAL80 substantially abrogated the majority of transgene expression mediated by *sNPF-R^R64H09^*, although a discrete population of cells in the ventral-lateral region of the antennal lobe (AL) retained expression in the presence of *elav-GAL80* (Fig. 2C-D). Consequently, the *sNPF-R^R64H09^* driver was found to label approximately 550 neuron-glia complexes, comprising 450-500 neurons and 50-100 glial cells (Fig. 2E-I).

**Figure 2.**
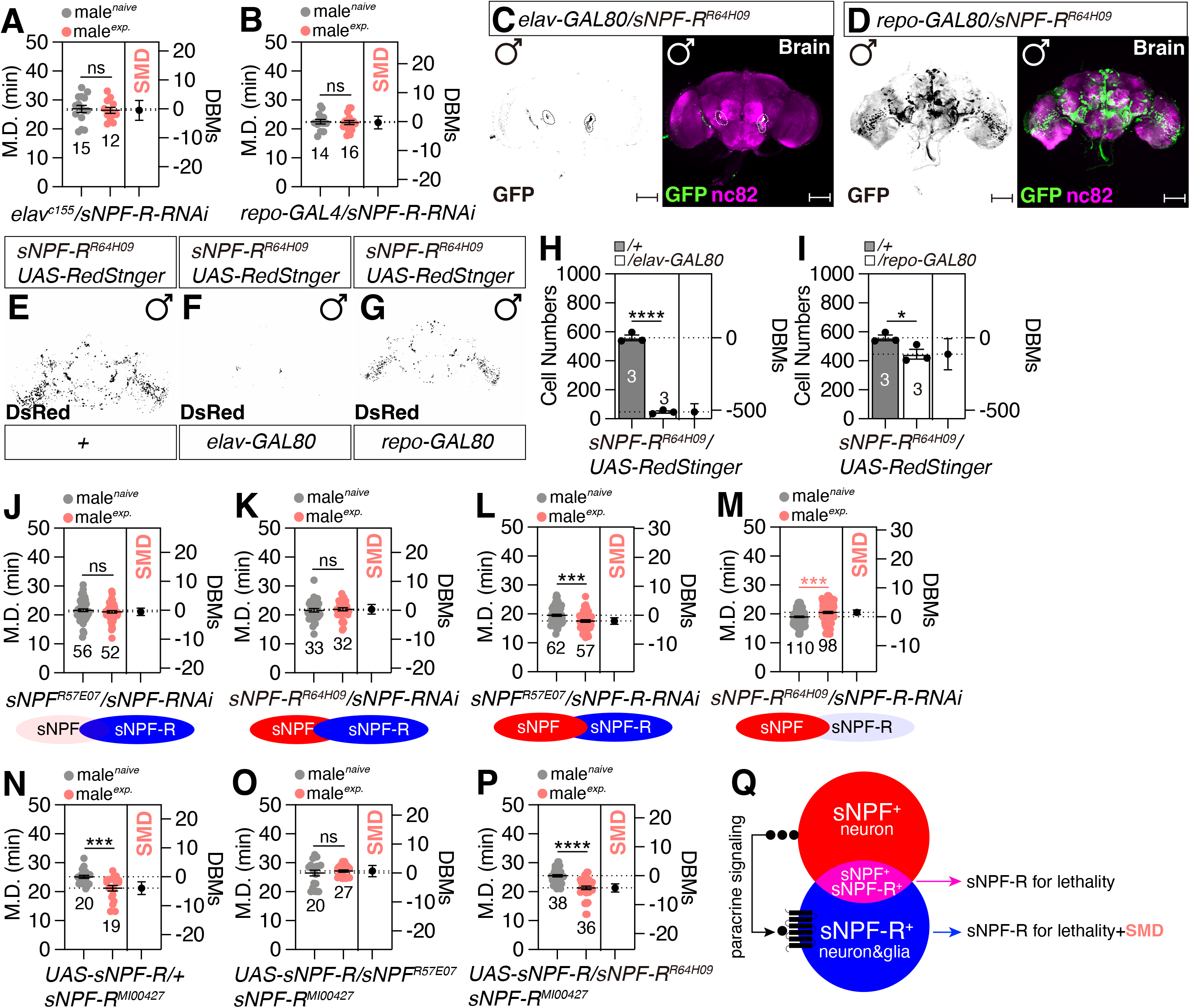
**sNPF-to-sNPF-R circuits are composed of functional neuron-glia network. (**A**-**B) SMD assay of *elav^c155^* and *repo-GAL4* mediated knockdown of sNPF-R *via sNPF-R-RNAi.* (C-D) Male flies expressing *elav-GAL80*, *sNPF-R^R64H09^* and *repo-GAL80*, *sNPF-R^R64H09^* with *UAS-mCD8GFP* were immunostained with anti-GFP (green), and anti-nc82 (magenta) antibodies. Scale bars represent 100 μm. (E-I) Brain of male flies expressing *sNPF-R^R64H09^* (G), *elav-GAL80*, *sNPF-R^R64H09^* (H) and *repo-GAL80*, *sNPF^R64H09^* (I) with *UAS-Redstinger* were immunostained with anti-DsRed (gray). Scale bars represent 100 μm. (H-I) Quantification of cell number. The flies are described as (E-G). Bars represent the mean *sNPF-R^R64H09^* (gray column) and *elav-GAL80*, *sNPF-R^R64H09^* (white column) cell number fluorescence level with error bars representing SEM (H). Bars represent the mean *sNPF-R^R64H09^* (gray column) and *repo-GAL80*, *sNPF-R^R64H09^* (white column) cell number fluorescence level with error bars representing SEM (I). (J-K) SMD assays for *sNPF^R57E07^* and *sNPF-R^R64H09^* mediated knockdown of sNPF *via* sNPF*-RNAi.* (L-M) *sNPF^R57E07^* and *sNPF-R^R64H09^* mediated knockdown of *sNPF-R via sNPF-R-RNAi.* (N-P) Genetic rescue experiments of SMD assays for overexpression of *sNPF-R via sNPF^R57E07^* and *sNPF-R^R64H09^* in *sNPF-R* mutant background flies. (Q) Diagram of connectivity between *sNPF* and *sNPF-R* in neuron and glia.

Subsequently, we sought to investigate the roles of sNPF and sNPF-R in cells co-expressing these two peptide and protein. We observed that the targeted suppression of sNPF in cells expressing either sNPF or sNPF-R led to a significant disruption in SMD behavior (Fig. 2J-K and Fig. S1F-G for genetic control). In contrast, the selective knockdown of sNPF-R in sNPF-R-expressing cells, but not in those expressing only sNPF, was sufficient to induce alterations in SMD (Fig. 2L-M and Fig. S1F-G for genetic control). These findings imply that sNPF plays an essential role in SMD within cells that co-express both components, whereas sNPF-R does not appear to be critical in those cells. Given that the *sNPF^R57E07^* driver is specific to neurons and not glia, our results suggest that sNPF primarily exerts its effects within neuronal cells. In contrast, sNPF-R seems to have a broader functional scope, mediating effects in both neuronal and glial cell populations.

Subsequently, our investigation aimed to elucidate the impact of varying local concentrations of sNPF and its receptor sNPF-R on the modulation of the SMD pathway. Overexpression of either sNPF or sNPF-R in cells positive for sNPF or sNPF-R did not affect SMD behavior (Fig. S1H-K and Fig. S1L-M for genetic controls), suggesting that the baseline expression levels of sNPF or sNPF-R in brain circuits are not critical for the regulation of SMD behavior.

The sNPF hypomorphic mutant flies exhibit viability (Lee et al. 2004; Lee et al. 2008), whereas the sNPF-R mutants consistently display a lethal phenotype (Enell 2011), indicating that the sNPF-R mutation exerts a more pronounced impact on *Drosophila* survival compared to the sNPF mutation. Utilizing a homozygous lethal mutant of sNPF-R^MI00427,^ we were able to rescue the lethal phenotype by expressing UAS-sNPF-R in cells positive for *sNPF^R57E0^* or *sNPF-R^R64H09^*. However, rescue of both the lethality and SMD behavior was only achieved when UAS-sNPF-R was expressed in sNPF-R-positive cells (Fig. 2N-P). These findings imply that sNPF-R expression in sNPF-positive neurons is not a prerequisite for the control of SMD behavior. In conclusion, our results indicate that the neuronal sNPF signaling to sNPF-R in either a neuronal or glial context, via a paracrine mechanism, is essential for the manifestation of SMD behavior (Fig. 2Q).

### A distinct subset of cells outside of mushroom body that express the sNPF-R is integral to the regulation of SMD behavior

The widespread expression of sNPF in a diverse population of neurons within the central nervous system of *Drosophila melanogaster* indicates that sNPF is not confined to a specific neuronal subtype or brain region but is expressed across various neuronal groups. This broad expression pattern suggests that sNPF may have multiple roles and functions within the CNS, potentially including the regulation of behaviors such as feeding, metabolism, stress responses, and reproduction. Consequently, identifying the specific functional roles of sNPF within each of these neuronal populations has proven to be a challenging endeavor (Nässel et al. 2008).

The mushroom body (MB) is a key structure in the *Drosophila* brain that is primarily associated with learning and memory (Modi et al. 2020). The expression of sNPF in the MB suggests that sNPF may play a role in regulating the processes of learning and memory. Several studies have indeed shown that sNPF and sNPF-R are involved in these memory functions in *Drosophila* (Johard et al. 2008; Knapek et al. 2013).

Nevertheless, the exact molecular and cellular mechanisms through which sNPF and its receptor sNPF-R modulate functions within the mushroom body remain to be elucidated.

The targeted knockdown of sNPF using the *GAL4^MB247^* driver did not alter SMD behavior; however, the knockdown of sNPF-R resulted in the disruption of SMD (Fig. 3A-B). In contrast, the knockdown of both sNPF and sNPF-R with the *GAL4^ok107^* driver significantly affected SMD, particularly resulting in sexually experienced males exhibiting an increased mating duration (indicated by pink asterisks in Fig. 3C-D). Neuron-specific knockdown of sNPF or sNPF-R, excluding *GAL4^MB247^*-positive MB neurons, yielded an SMD phenotype identical to that observed following *GAL4^ok107^*-mediated knockdown of sNPF or sNPF-R (compare Fig. 3C-D to 3E-F, Fig. S2A-C for genetic control).

**Figure 3.**
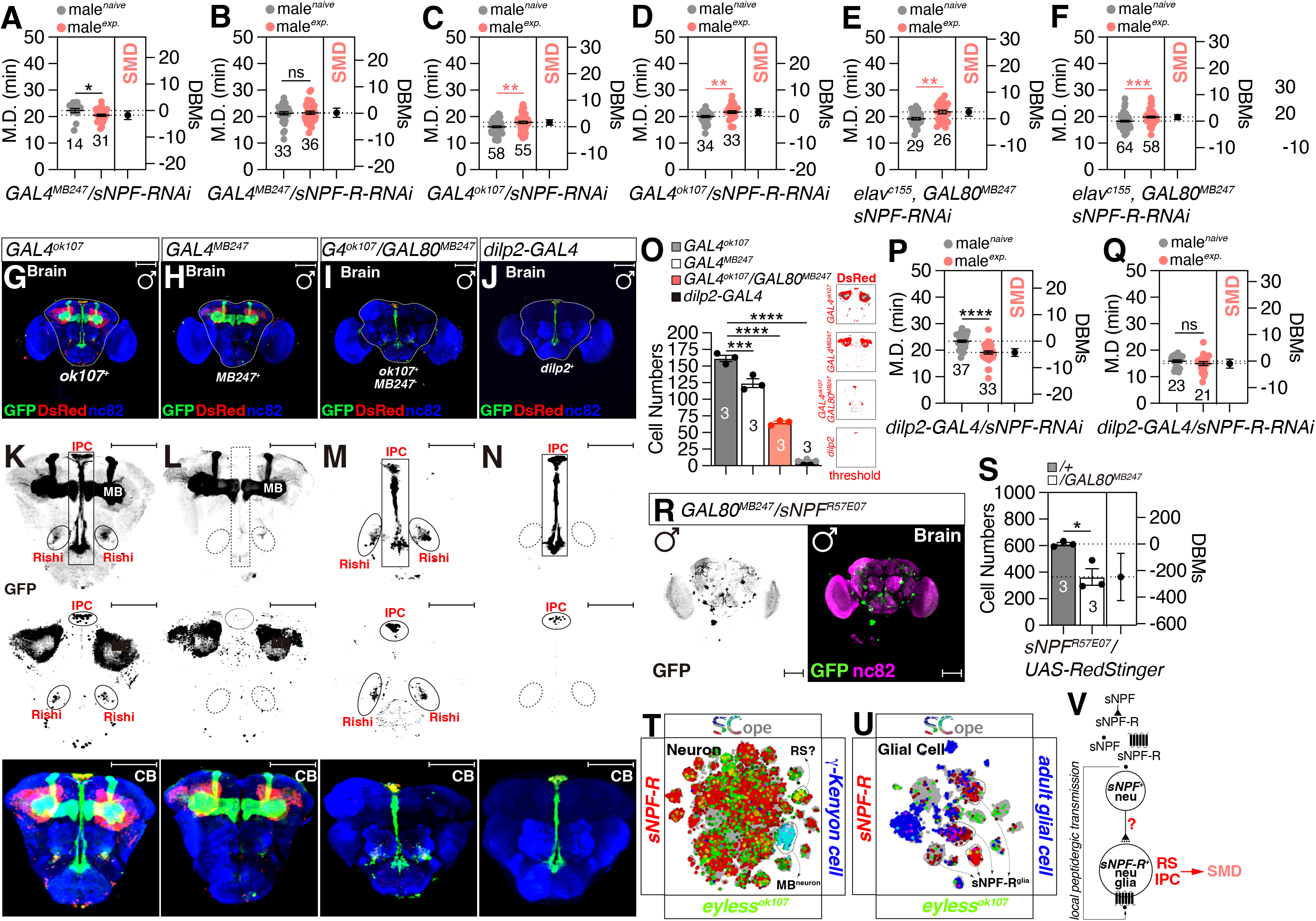
A distinct subset of cells outside of mushroom body that express the sNPF-R is integral to the regulation of SMD behavior. (A-D) SMD assays of mushroom driver *GAL4^MB247^*and *GAL4^ok107^* mediated knockdown of *sNPF* and *sNPF-R via sNPF-RNAi* and *sNPF-R-RNAi*. (E-F) SMD assays of *elav^c155^* mediated knockdown of *sNPF* and *sNPF-R via sNPF-RNAi* together with *GAL80^MB247^* and *sNPF-R-RNAi* together with *GAL80^MB247^*. (G-I) Brain of male flies expressing *GAL4^MB247^* (G), *GAL4^ok107^* (H) and *GAL4^ok107^*, *GAL80^MB247^* (I) together with *UAS-CD8tdGFP, UAS-Redstinger* were immunostained with anti-GFP (green), anti-DsRed (red) and nc82 (blue) antibodies. Scale bars represent 100 μm. (J) Brain of male flies expressing *dilp2-GAL4* together with *UAS-CD8tdGFP, UAS-Redstinger* were immunostained with anti-GFP (green), anti-DsRed (red) and nc82 (blue) antibodies. Scale bars represent 100 μm. (K-N) Central brain (CB) of male flies are described as (G-J). Scale bars represent 100 μm. (O) Quantification of cell number. The flies are described as (G-N). Bars represent the mean *GAL4^ok107^* (black column), *GAL4^MB247^* (white column), the combination *GAL4^ok107^*, *GAL80^MB247^* (red column) and *dilp2-GAL4* (gray column) cell number fluorescence level with error bars representing SEM. (P-Q) SMD assays of *dilp2-GAL4* mediated knockdown of *sNPF* and *sNPF-R via sNPF-RNAi* and *sNPF-R-RNAi*. (R-S) Brain of male flies expressing *GAL80^MB247^* and *sNPF-R^R57E07^* together with *UAS-CD8tdGFP* were immunostained with anti-GFP (green) and nc82 (magenta) antibodies. Scale bars represent 100 μm. Bars represent the mean *sNPF^R57E07^* (gray column) and *sNPF^R57E07^*, *GAL80^MB247^* (white column) cell number fluorescence level with error bars representing SEM. (T-U) Single-cell RNA sequencing (SCOPE scRNA-seq) datasets reveal cell clusters colored by expression of *sNPF-R* (red), *eyeless^ok107^ /*γ*-Kenyon cell* (green/blue) in neuron (T), *sNPF-R* (red), *eyeless^ok107^ /*adult glial c*ell* (green/blue) in glial cell (U). (V) Diagram of connectivity between *sNPF* and *sNPF-R*.

The *GAL4^MB247^* and *GAL4^ok107^* drivers are both commonly used in fruit fly research to label and manipulate specific subsets of MB neurons. *GAL4^MB247^* drives expression in a subset of Kenyon cells. These cells are primarily located in the α/β lobes of the MB. *GAL4^ok107^*, on the other hand, labels the most of MB neurons as well as additional regions outside the MB (Aso et al. 2009; Kaun et al. 2011).

Due to the absence of comprehensive expression analysis, we conducted a comparative examination of the expression patterns of *GAL4^ok107^* (Fig. 3G and K) and *GAL4^MB247^* (Fig. 3H and L), revealing that *GAL4^ok107^*encompasses additional brain regions, including IPCs in the pars intercerebralis (PI) and an as yet unidentified population of neurons adjacent to the ventral-lateral region of the AL (Fig. 3I-J and 3M-N). Lacking prior information on the specific identity of these cellular clusters, we have preliminarily designated them as ’Rishi’ cells (RS). This designation is based on the cells’ morphology, which is reminiscent of a solar eclipse. The term ’Rishi’ is derived from Chinese, meaning solar eclipse, and is chosen to reflect the dual nature of these cells, which, like the sun and moon in an eclipse, represent a functional interaction between neurons and glia.

Quantification of cell numbers revealed approximately 60 *GAL4^ok107^*-positive, *GAL4^MB247^*-negative cells, with around 10 of these being dilp2-positive, suggesting the presence of approximately 50 RS cells in the male brain (Fig. 3O). The membrane-covered area correlated with the number of cells labeled by each GAL4 driver (Fig. S2D). Knockdown of sNPF in IPCs did not alter SMD behavior (Fig. 3P), while knockdown of sNPF-R in these cells did disrupt SMD (Fig. 3Q and Fig. S2E for genetic control), indicating that sNPF-R expression in regions outside the MB, including IPCs and RS cells, is critical for the regulation of SMD behavior.

Co-expression with *GAL80^MB247^* resulted in the disappearance of *sNPF^R57E07^*- and *sNPF-R^R64H09^*-positive MB projections (Fig. 3R), and approximately 250 and 150 cells among the sNPF- and sNPF-R-positive populations were identified as *GAL4^MB247^*-positive (Fig. S2F-J and Fig. 3S). Analysis of Fly SCope data suggests the presence of sNPF-R-positive and eyeless-positive (gene for *GAL4^ok107^*) neurons and glial cells both within and outside the MB (Fig. 3T-U). Utilizing Virtual Fly Brain (VFB), an interactive tool for *Drosophila* neuroanatomy and gene expression (Court et al. 2023), in conjunction with recently developed subtype glia-specific Fly Light GAL4 drivers (Pfeiffer et al. 2008; Malte C. Kremer et al. 2017), we determined ALG and SPG overlap with the RS cells labeled by the *sNPF-R^R64H09^*driver (Fig. S2K).

Examination through the 3D viewer confirmed that both ALG and SPG are associated with the anterior RS cells, with only ALG overlapping with the posterior RS cells (Movie. 1).

Collectively, these findings imply that paracrine signaling mediated by neuronal sNPF emanating from outside the MB to IPC or RS cells expressing sNPF-R is a pivotal mechanism underlying the emergence of SMD behavior (Fig. 3V).

### The astrocyte-like glial expression of sNPF-R is essential for the manifestation of SMD behavior

Both vertebrate and *Drosophila* glia have similar fundamental functions and anatomical structures (Freeman and Doherty 2006). The neural systems of adult fruit flies consist of six different types of glia. The cortical regions contain cortex glia (CG), while the neuropile regions contain astrocyte-like (ALG). In the peripheral nervous system (PNS), there are wrapping glia associated with central axon tracts (EGT) and peripheral nerves (EGN). Additionally, there are two sheet-like glia, subperineurial (SPG) and perineurial (PNG), which form a surface that covers both neuronal systems (Malte C. Kremer et al. 2017). To delineate the specific subtypes of glial cells that express sNPF-R and contribute to the generation of SMD behavior, we employed recently characterized glial subtype-specific GAL4 drivers (Malte C Kremer et al. 2017). Our findings indicated that the expression of sNPF-R in ALG, CG, EGT, and SPG is necessary for the manifestation of SMD behavior (Fig. 4A-F and Fig. S3A-F for genetic control). In particular, the reduction of sNPF-R expression in ALG neurons results in sexually experienced male flies exhibiting prolonged mating durations compared to inexperienced counterparts (Fig. 4A). This observation implies that ALG neurons may represent a distinct glial subtype that is crucial for the sNPF-R’s role in modulating SMD behavior.

**Figure 4.**
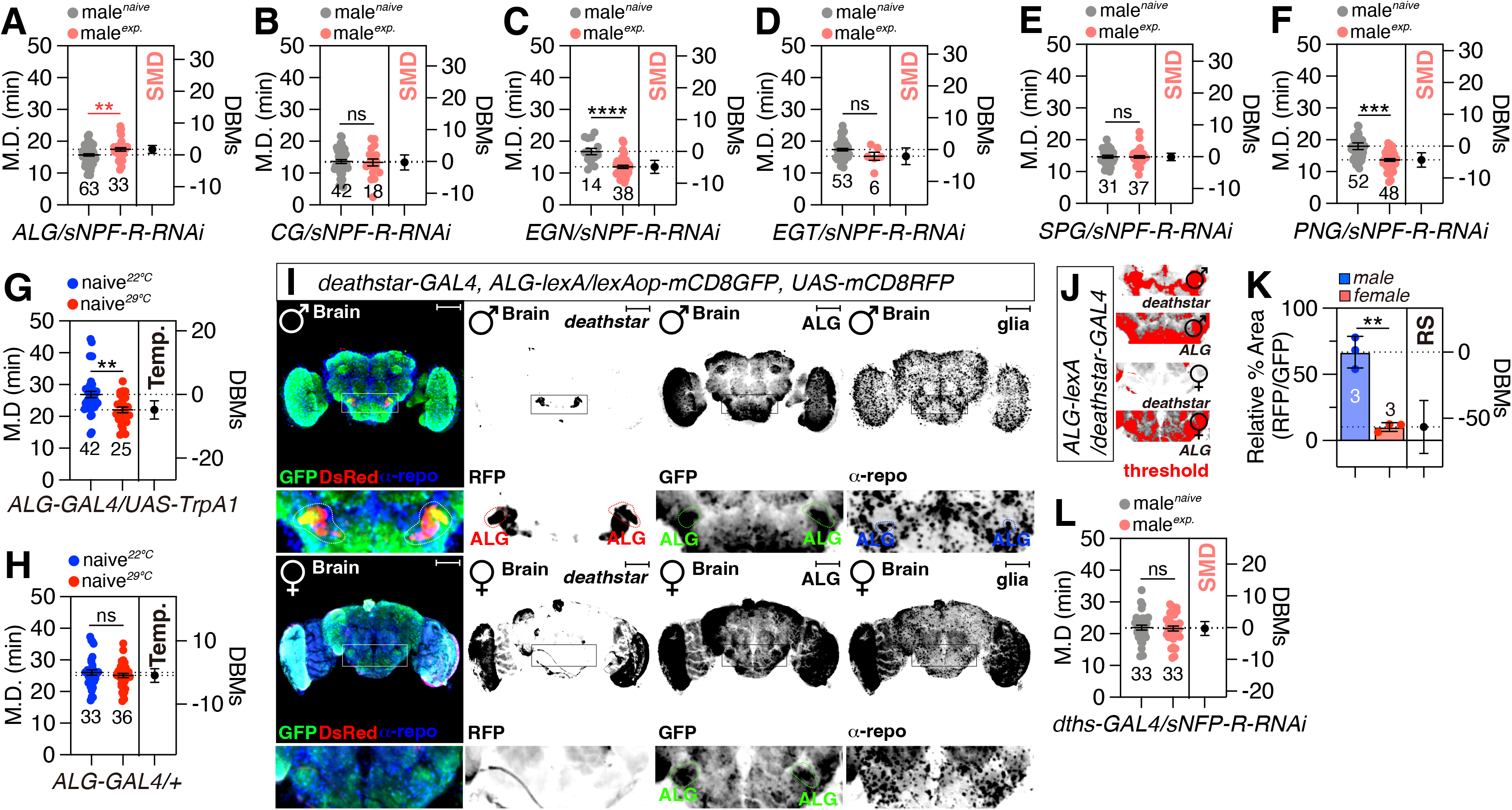
The astrocyte-like glial expression of sNPF-R is essential for the manifestation of SMD behavior. (A-F) SMD assays for *ALG-GAL4* (A)*, CG-GAL4* (B)*, EGN-GAL4* (C)*, EGT-GAL4* (D)*, SPG-GAL4* (E)*, PNG-GAL4* (F), mediated knockdown of *sNPF-R via sNPF-R-RNAi.* (G-H) TMD assay for active *sNPF^R57E07^* with *ALG-GAL4* (G)*, ALG-GAL4/+* at 29[. (I) Co-localization analysis of *deathstar-GAL4* and *ALG-lexA* with *UAS-mCD8GFP, UAS-mCD8RFP* were immunostained with anti-GFP (green), anti-DsRed (red) and nc82 (magenta) antibodies. The yellow area represents overlap by *deathstar-GAL4*(red) and *ALG-lexA* (green). (J-K) Quantification of the relative percent area formed by *deathstar-GAL4* (red) and *ALG-lexA* (green) in the brains of male and female flies was conducted. (L) SMD assays for *deathstar-GAL4* mediated knockdown of *sNPF-R via sNPF-R-RNAi.*

It is well-established that sNPF acts as an inhibitory modulator within a diverse array of neuronal circuits (Lee et al. 2008; Shang et al. 2013; Vecsey et al. 2014). To elucidate the specific roles of sNPF signaling in various glial subtypes, we utilized a constitutively active form of the bacterial sodium channel *NaChBac* to artificially perturb the electrical properties of glial cells. By introducing *NaChBac* into distinct glial populations, we induced hyperexcitation, a strategy previously demonstrated to disrupt ionic gradients and potentially impact transporter functions in glial cells (Nitabach et al. 2006; Beart and O’Shea 2007; Charalambous and Wallace 2011).

*NaChBac* expression in glial cells of *Drosophila* has been shown to disrupt circadian rhythms, suggesting a physiological influence on circadian pacemaker neurons (Ng et al. 2011). Our results indicate that the activation of ALG and EGT glial cells specifically disrupts SMD behavior (Fig. S3G-L and Fig. S3M for genetic control), suggesting that these glial populations are critical for sNPF-R-mediated SMD behavior.

To further delineate the contributions of individual glial subtypes to SMD behavior, we activated glial cells using the heat-sensitive *Drosophila* cation channel TrpA1 and transferred the experimental flies to the activation temperature of 29°C (Hamada et al. 2008; Kang et al. 2010; Du et al. 2016). Among the tested glial populations, activation of ALG cells led to a significant reduction in mating duration (Fig. 4G and Fig. 4H for genetic control), indicating that ALG might be the most critical glial population for sNPF-sNPF-R circuit function in energy balance regulation. Consequently, the equilibrium between the inhibitory influence of sNPF-R on ALG neurons and their electrical excitability is pivotal for the manifestation of SMD behavior, which is essential for maintaining energy homeostasis following mating experience.

We confirmed this hypothesis using a newly identified *ALG*-specific gene *deathstar* (*dths*) driver strain, *dths-GAL4* (X. Zhang et al. 2024), which revealed that ALG cells located near the ventral-lateral region of the AL, similar to the location of RS cells, are pivotal for sNPF-sNPF-R signaling in energy balance regulation (Fig. 4I).

Interestingly, neurons labeled by *dths-GAL4* displayed sexual dimorphism, which was observed exclusively in the male brain (compare male and female panels in Fig. 4I), indicating that these neurons may specifically contribute to male physiological processes (Fig. 4J-K for quantification). Knockdown of sNPF-R using *dths-GAL4* disrupted SMD behavior (Fig. 4L and Fig. S3N for genetic control), suggesting that sNPF-R-expressing ALG cells can be labeled by *dths-GAL4*. We have recently identified that the *dths* gene specifically modulates adult locomotion behavior (Najafi et al. 2022). Knockdown of *dths* using the *sNPF-R^R64H09^*driver resulted in a more pronounced reduction in locomotion behavior in males compared to females (Fig. S3O-P), suggesting that *dths* expression in ALG plays a role in modulating SMD behavior. These data collectively suggest that glial function within sNPF-R expressing cells, particularly ALG, plays a critical role in regulating SMD behavior within the sNPF-sNPF-R circuit (Fig. S3Q).

### The sexually dimorphic nature of sNPF-sNPF-R neural circuits constitutes a pivotal regulatory mechanism for SMD behavior

Sexual dimorphism is the difference in structure or function between male and female of the same species. Sexual dimorphism in the brain of the fruit fly can affect neuroanatomy, neurotransmitter levels, and gene expression (Yamamoto et al. 1998).

In our investigation, we discovered sexual dimorphism in the expression of *sNPF-R^R64H09^*-positive cells within the RS cells. Quantitative analysis revealed that the male brain harbors approximately 600 sNPF-R-positive cells, whereas the female brain contains only 400 such cells. Additionally, male sNPF-R-positive cells occupy a broader anatomical area of the brain compared to females, while the size of these cells remains comparable between the sexes (Fig. 5A-E). The sexually dimorphic expression pattern of sNPF-R was consistent across the central brain (CB) and OL (Fig. 5F-H). Feminizing these cells resulted in a reduction in cell number and brain coverage, without altering cell size (Fig. 5I-L and Fig. S4A).

**Figure 5.**
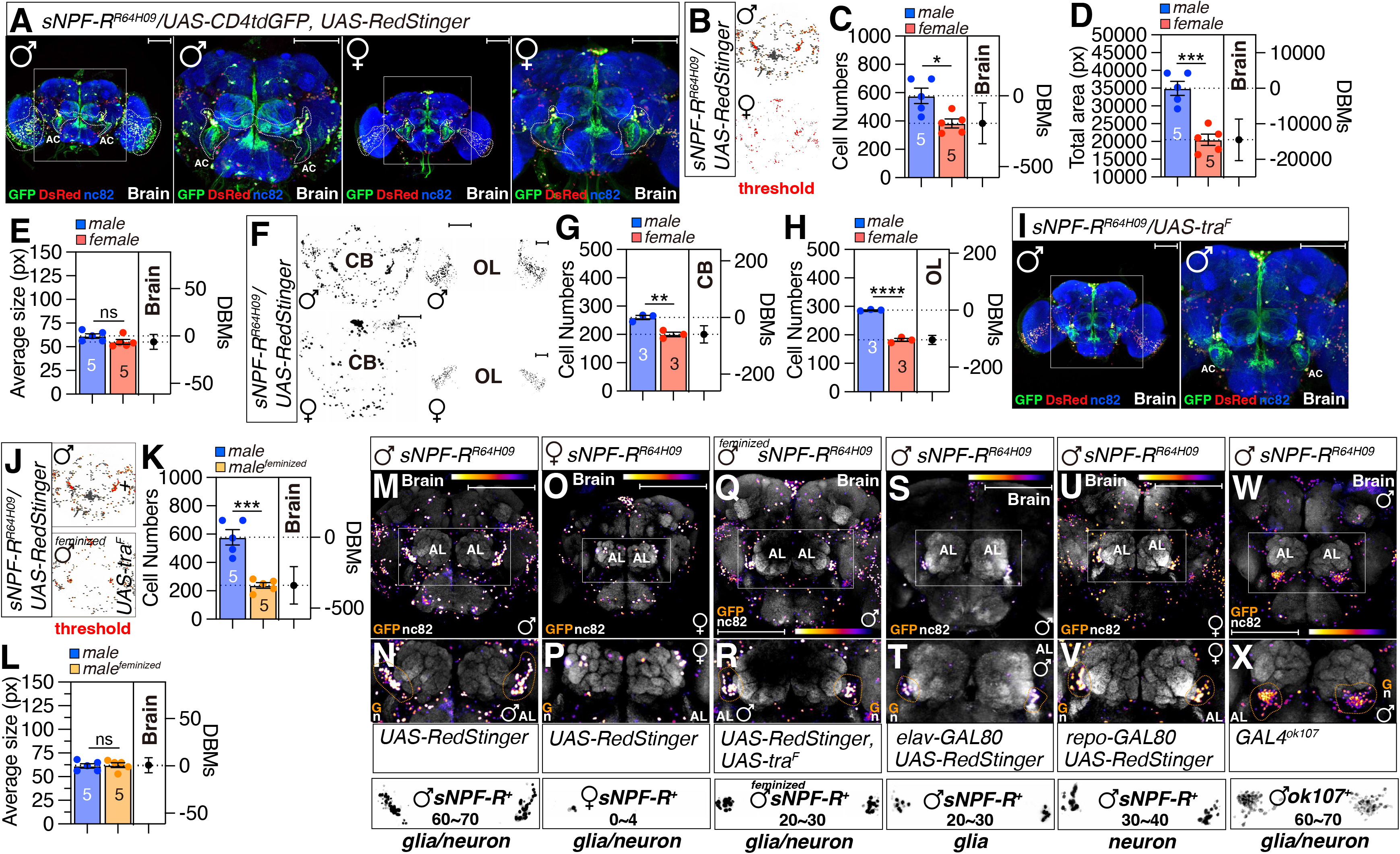
The sexually dimorphic nature of sNPF-sNPF-R neural circuits constitutes a pivotal regulatory mechanism for SMD behavior. (A) Brain of male and female flies expressing *sNPF-R^R64H09^* together with *UAS-CD4tdGFP, UAS-Redstinger* were immunostained with anti-GFP (green), anti-DsRed (red) and nc82 (blue) antibodies. Scale bars represent 100 μm. Boxes indicate the magnified regions of interest presented in the middle panels. Scale bars represent 100 μm. Boxes indicate the magnified regions of interest presented in the middle panels. (B) The RFP fluorescence in male and female fly brains expressing *sNPF-R^R64H09^* with *UAS-Redstinger* was processed using ImageJ software, a threshold function was applied to distinguish fluorescence from the background. (C-E) Quantification of *sNPF-R^R64H09^* cell number (C), cell relative area (D), and cell average size (E) between males (blue column) and females (red column) showing fluorescence levels with error bars representing SEM. (F) The RFP fluorescence in the central brain (CB) and optic lobe (OL) of male and female flies expressing *sNPF-R^R64H09^* with UAS-Redstinger was processed using ImageJ software. A threshold function was applied to distinguish fluorescence from the background. (G-H) Quantification of *sNPF-R^R64H09^* cell number between males (blue column) and females (red column) showing fluorescence levels with error bars representing SEM. (I) Brain of male expressing *sNPF-R^R64H09^*, *UAS-traF* with *UAS-CD4tdGFP, UAS-Redstinger* were immunostained with anti-GFP (green), anti-DsRed (red) and nc82 (blue) antibodies. Scale bars represent 100 μm. Boxes indicate the magnified regions of interest presented in the middle panels. (J-L) The RFP fluorescence in male and feminized male fly brains expressing *sNPF-R^R64H09^*, *UAS-traF* with *UAS-Redstinger* was processed using ImageJ software, a threshold function was applied to distinguish fluorescence from the background (J). Quantification of *sNPF-R^R64H09^* cell numbers (K) and cell average size (L) between males (blue column) and feminized males (yellow column), displaying fluorescence levels with error bars representing SEM. (M-X) Brains of male and female flies (M, O) expressing *sNPF-R^R64H09^* with *UAS-Redstinger* and brains of male flies (Q, S, U, W) expressing *sNPF-R^R64H09^*, *UAS-traF*(Q), *sNPF-R^R64H09^*, *elav-GAL80*(S), and *sNPF-R^R64H09^*, *repo-GAL80* (U), *GAL4^ok107^*, *GAL80^MB247^* (W) were immunostained with anti-DsRed (fire) and nc82 (gray) antibodies. Boxes (N, P, R, T, V) indicate the magnified regions of interest and cell numbers near the AL regions.

Despite the well-documented sexual dimorphism in *Drosophila*, the sexually dimorphic functions of sNPF have not been previously explored. Our data indicate a clear sexual dimorphism in the expression of *sNPF^R57E07^*, with males exhibiting a greater number of cells and broader brain coverage, while maintaining a similar cell size compared to females (Fig. S4B-F). These findings suggest that sNPF expression is also subject to sexual dimorphism, with males possessing a higher density of sNPF-positive neurons.

Through a comparative analysis of transgene combinations, we determined that the male brain contains 60-70 sNPF-R-positive glia/neuron complexes in the RS cells, of which 20-30 are glial cells and 30-40 are neurons, a distribution similar to that observed in *GAL4^ok107^*-positive RS cells (Fig. 5M-T). In contrast, females and feminized males exhibit a reduced number of RS cells, ranging from 20-30 (Fig. 5O-X). This evidence strongly suggests that sNPF-R-expressing RS cells are sexually dimorphic, with a higher representation in the male brain.

The transcription factors *fruitless* (*fru*) and *doublesex* (*dsx*) are integral to the establishment of sexual dimorphism in *Drosophila*. *Fru* is predominantly active in the male nervous system and is essential for the development of male-specific courtship behaviors. *Dsx*, on the other hand, is pivotal in directing somatic cells towards male- or female-specific differentiation. The interplay between these genes and the intricate regulatory networks in which they operate underlies the coordination of sexual differentiation in *Drosophila* (Yamamoto et al. 1998; Pan and Baker 2014; Zhou et al. 2014).

Our findings reveal that *sNPF-R^R64H09^* labels a subset of fru-positive cells in the RS cells (Fig. 6A), while *sNPF^R57E07^* identifies a substantial number of fru-positive neurons in the OL and CB (Fig. S4G). Approximately 50 sNPF- and fru-positive neurons are present in the CB, and 160 in the OL (Fig. S4H). Inhibition of these neurons through the expression of tetanus toxin light chain (TNT) or ricin, as well as feminization of sNPF neurons, disrupts SMD behavior (Fig. 6B-C and Fig. S5A).

**Figure 6.**
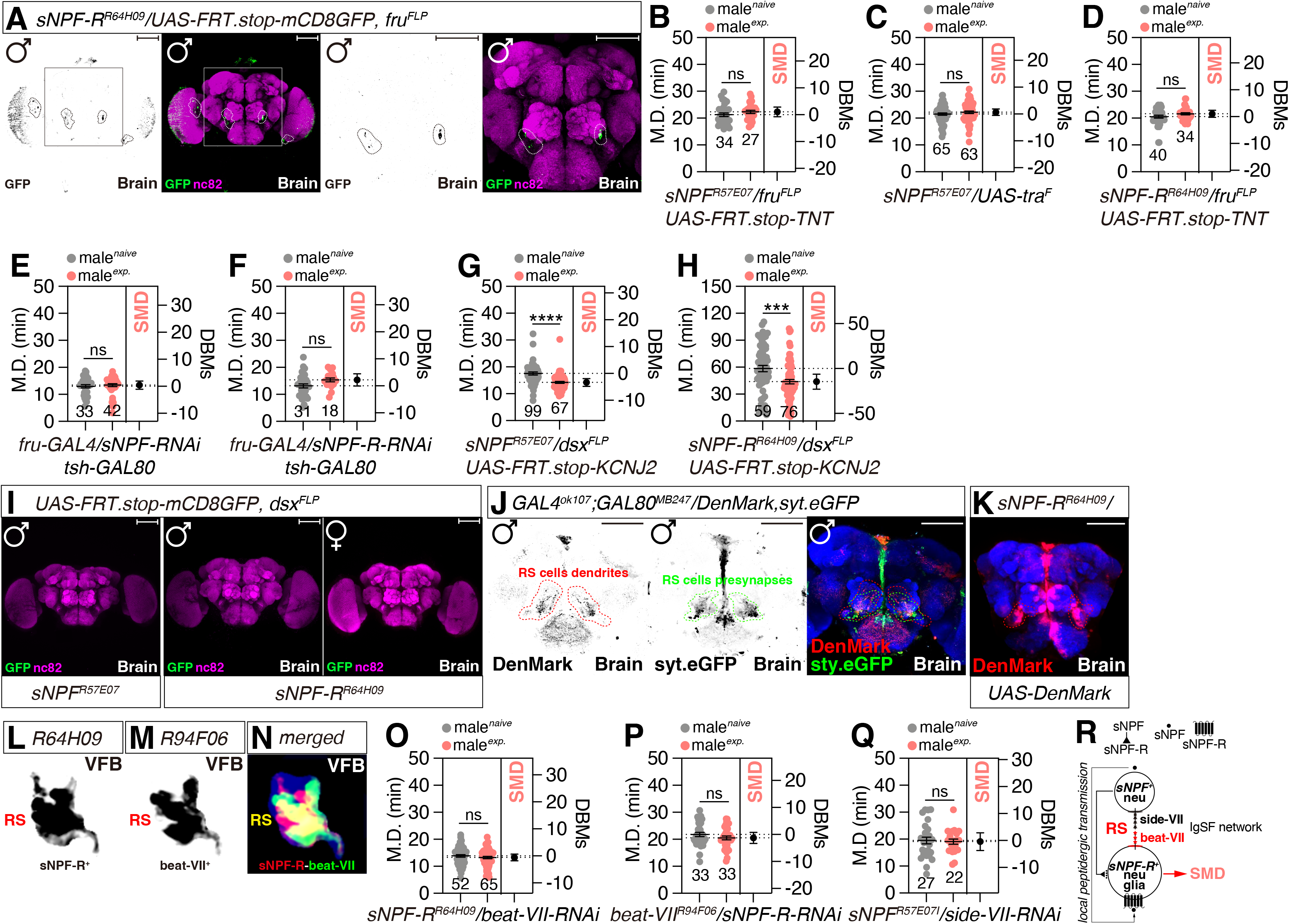
Neuronal anatomy and structural features of RS cells. (A) Brain of male flies expressing *sNPF-R^R64H09^* with *UAS-FRT.stop-mCD8GFP, fru^FLP^* were immunostained with anti-GFP (green), and nc82 (magenta) antibodies. Scale bars represent 100 μm. Boxes indicate the magnified regions of interest presented in the middle panels. (B-C) SMD assays of *sNPF^R57E07^* cross with *UAS-FRT.stop-TNT, fru^FLP^* and *UAS-tra^F^.* (D) SMD assays of *sNPF-R^R64H09^* cross with *UAS-FRT.stop-TNT, fru^FLP^.* (E-F) SMD assays of *fru-GAL4, tsh-GAL80* mediated knockdown of *sNPF* and *sNPF-R via sNPF-RNAi* and *sNPF-R-RNAi*. (G-H) SMD assays of *sNPF^R57E07^* and *sNPF-R^R64H09^* cross with *UAS-FRT.stop-KCNJ2, dsx^FLP^.* (I) Brain of male flies expressing *sNPF^R57E07^* and *sNPF-R^R64H09^*with *UAS-FRT.stop-mCD8GFP, dsx^FLP^* were immunostained with anti-GFP (green), and nc82 (magenta) antibodies. Scale bars represent 100 μm. (J) Brain of male flies expressing *GAL4^ok107^*, *GAL80^MB247^* with *UAS-syteGFP, UAS-denmark* were immunostained with anti-GFP (green), anti-DsRed (red) and nc82 (blue) antibodies. Scale bars represent 100 μm. (K) Brain of male flies expressing *sNPF-R^R64H09^*with *UAS-denmark* were immunostained with anti-DsRed (red) and nc82 (blue) antibodies. Scale bars represent 100 μm. (L-M) Expression patterns of RS region for *sNPF-R^R64H09^* (L) and *beat-VII ^94F06^* (M) in the Virtual Fly Brain (VFB). (N) Co-localization analysis of *beat-VII ^94F06^* and *sNPF-R^R64H09^* in Virtual Fly Brain (VFB). The yellow area represents overlap by *beat-VII ^94F06^*(green) and *sNPF-R^R64H09^* (red). (O) SMD assays for *sNPF-R^R64H09^*mediated knockdown of *beat-VII* via *beat-VII-RNAi.* (P) SMD assays for *beat-VII-GAL4* mediated knockdown of *sNPF-R* via *sNPF-R-RNAi.* (Q) SMD assays for *sNPF^R57E07^* mediated knockdown of *side-VII* via *side-VII-RNAi.* (R) Diagram the network involving sNPF, sNPF-R, beat-VII, and side-VII in neurons and glia.

However, the expression of an inactive form of TNT did not affect SMD behavior (Fig. S5B). For the genetic control experiments, please consult the relevant literature (Lee et al. 2023).

Similarly, inhibition or ablation of sNPF-R-positive and fru-positive cells using TNT or ricin disrupts SMD behavior (Fig. 6D, Fig. S5C), while the inactive form of TNT does not alter energy balance behavior (Fig. S5D). Feminization of *sNPF-R^R64H09^* cells results in defects across the spectrum of male mating behaviors (Fig. S5E), indicating that fru-positive RS cells are critical for the execution of most male sexual behaviors. Knockdown of sNPF or sNPF-R in fru- or dsx-positive brain cells also disrupts SMD behavior (Fig. 6E-F and Fig. S5F), but inhibition of dsx-positive *sNPF^R57E07^* neurons (Fig. 6G) or dsx-positive and *sNPF-R^R64H09^* neurons (Fig. 6H) does not affect SMD behavior. As anticipated, we did not observe any dsx-positive *sNPF^R57E07^* or *sNPF-R^R4H09^* cells in either the male or female brain (Fig. 6I).

### Neuronal anatomy and structural features of RS cells

To delineate the neuronal architecture of the RS cells in male fly brain, we employed the UAS-*Denmark*, *syt.eGFP* marker to trace the dendritic arbors and presynaptic terminals of RS cells (Nicolaï et al. 2010). *GAL4^MB247^*failed to yield any dendritic or presynaptic labeling in the vicinity of the RS cells (Fig. S5G). In contrast, *GAL4^ok107^* robustly labeled both dendritic and presynaptic structures adjacent to the RS cells (Fig. S5H), even after the exclusion of *GAL4^MB247^*-positive Kenyon cells (Fig. S5I). The dendritic signals emanating from *GAL4^ok107^*-positive yet *GAL4^MB247^*-negative cells were observed across the medial and lateral inferior regions, while the presynaptic neuronal processes predominantly projected to locales proximate to the RS cells (Fig. 6J). The *sNPF-R^R64H09^* driver recapitulated a similar neuroanatomical profile to that observed with *GAL4^ok107^*, indicating that RS cells are likely to engage in bidirectional signaling with local sNPF neurons (Fig. 6K). This anatomical distribution suggests that these RS cells receive inputs from medial and lateral inferior brain regions, as well as from local sNPF peptidergic transmission zones.

Virtual Fly Brain (VFB) 3D analysis further confirmed that *sNPF^R57E07^* and *sNPF-R^R64H09^* cells extend processes to both lateral-inferior and RS cell region markers and form synapses with each other (Fig. S5J-L). Utilizing VFB connectome data, we identified that the GAL4 driver *beat-VII^R94F06^* also targets RS cells (Fig. S5M) and exhibits colocalization with *sNPF-R^R64H09^* within the RS cell region (Fig. S5N-S). The beat-VII (beaten path VII) gene in *Drosophila* encodes an immunoglobulin superfamily (IgSF) protein predicted to mediate heterophilic cell-cell adhesion via plasma membrane interactions and is expressed in the adult head (Fambrough and Goodman 1996; Pipes et al. 2001; Vogel et al. 2003; Tan et al. 2015). The beat family proteins contribute to the fly cell surface interactome, with Sidestep (side) serving as a ligand-receptor pair critical for axon guidance (Li et al. 2017).

The *GAL4^R94F06^*driver of the beat-VII promoter revealed clear overlap with *sNPF-R^R64H09^* cells within the RS cells (Fig. 6L-N). Knockdown of beat-VII in *sNPF-R^R64H09^*-positive cells or knockdown of sNPF-R in *beat-VII^R94F06^*-positive cells resulted in the abolition of SMD behavior in male flies (Fig. 6O-P), akin to the effects observed upon knockdown of side-VII, potential receptors for beat-VII-mediated cell-cell interactions (Li et al. 2017; Osaka et al. 2023) (Fig. 6Q). These findings suggest that RS cells represent a specialized feature of sNPF-R expressing cells, which rely on the Beat-Side interactome to ensure precise local peptidergic transmission between sNPF and sNPF-R (Fig. 6R).

### Sexual experience induces global alterations in calcium signaling throughout sNPF-sNPF-R brain circuits

Male sexual experience triggers widespread alterations in calcium dynamics across the brain-body axis, manifesting as a shift in internal states facilitated by SIFa-SIFaR signaling pathways (Wong et al. 2019). Sexual experience elicits a pronounced increase in calcium levels within *GAL4^ok107^*-labeled MB, PI, AL, and SOG (Fig. 7A-G, Fig. S6A-D). Notably, the calcium levels in the dorsal accessory calyx (dAC), a region renowned for its multisensory integration of olfactory, gustatory, and visual inputs (Aso et al. 2014), undergo a significant surge following male’s sexual experience (Fig. 7H). Furthermore, sexual experience precipitates marked modifications in *sNPF-R^R64H09^*-expressing cells in the brain (Fig. S6E-F), particularly within the SOG and AL domains where RS cells are situated (Fig. 7I-M), while the OL remain unaffected (Fig. 7N-O). These data suggest that alterations in global calcium signaling within the sNPF-sNPF-R circuit, including the MB dAC and RS cells, are associated with male sexual experience to regulate mating duration.

**Figure 7.**
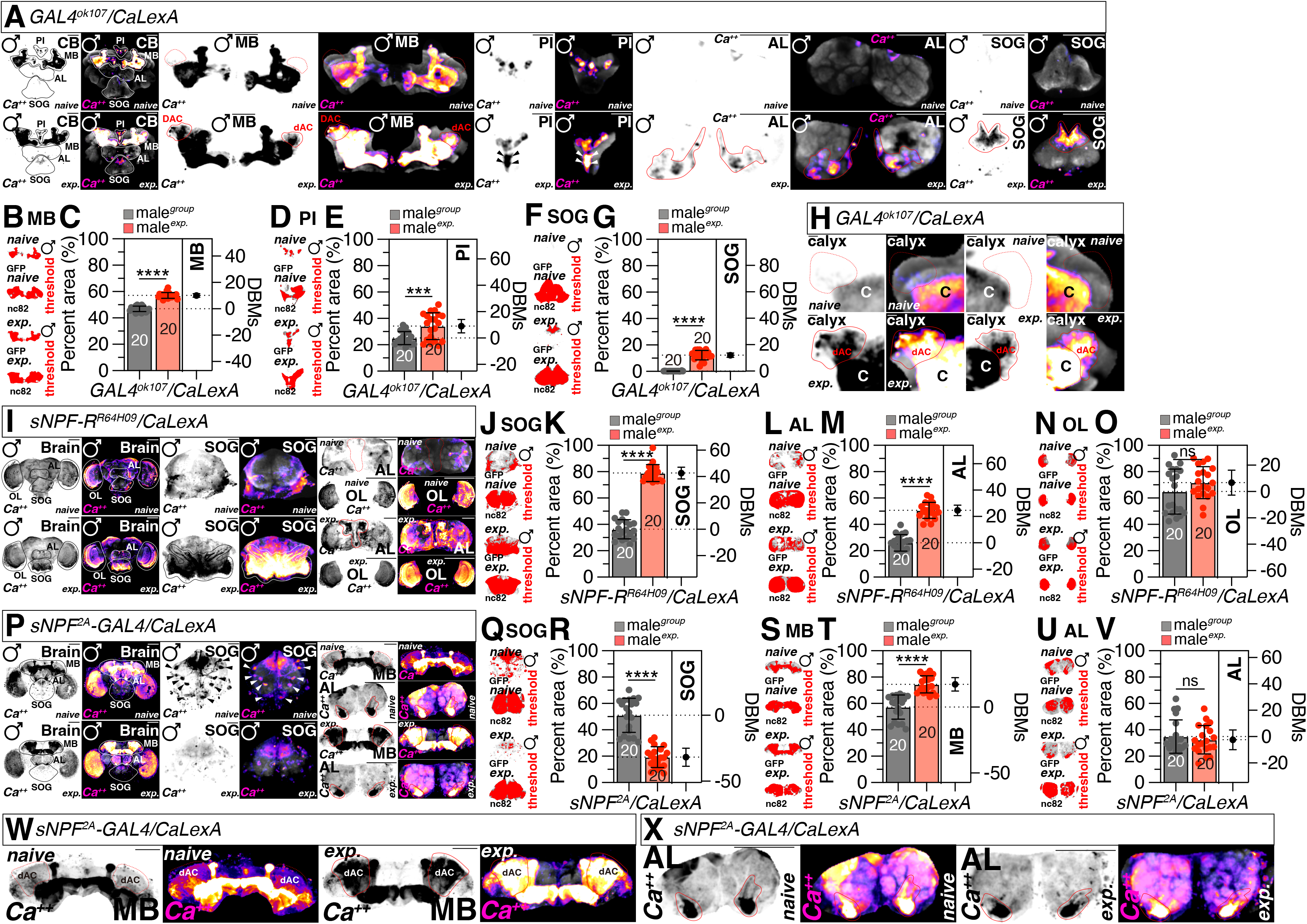
Sexual experience induces global alterations in calcium signaling throughout sNPF-sNPF-R brain circuits. (A) Different levels of neural activity of the brain as revealed by the CaLexA system in naïve and experienced flies. Male flies expressing *GAL4^ok107^* along with *LexAop-CD2-GFP, UAS-mLexA-VP16-NFAT and LexAop-CD8-GFP-A2-CD8-GFP* were dissected after 5 days of growth (mated male flies had 1 day of sexual experience with virgin females). The dissected brains were then immunostained with anti-GFP (green) and anti-nc82 (blue). GFP is pseudo-colored as “red hot”. Boxes indicate the magnified regions of interest presented in the right panels. Scale bars represent 100 μm. (B-G) Quantification of GFP fluorescence in MB (B-C), PI (D-E), SOG (F-G) between naïve and experienced *GAL4^ok107^* male flies. The GFP fluorescence was processed using ImageJ software, applying a threshold function to distinguish fluorescence from the background (B, D, F) and normalizing the GFP fluorescence relative to the nc82 fluorescence. The conditions of flies are described above: naïve, naïve male flies; exp., male flies with sexual experience. Bars represent the mean of the normalized GFP fluorescence level with error bars representing the SEM. Asterisks represent significant differences, as revealed by the Student’s *t* test and ns represents non-significant difference (*\*p < 0.05, **p < 0.01, ***p < 0.001*). See the STAR 17 METHODS for a detailed description of the fluorescence intensity analysis used in this study. (H) Different levels of neural activity in the dAC region of male flies expressing *GAL4^ok107^* along with the CaLexA system were observed between naïve and experienced flies. (I) Different levels of neural activity in the SOG and OL regions of male flies expressing *sNPF-R^R64H09^* along with the CaLexA system were observed between naïve and experienced flies. (J-O) Quantification of GFP fluorescence in SOG (J-K), AL (L-M), OL (N-O) between naïve and experienced *sNPF-R^R64H09^* male flies. The GFP fluorescence was processed using ImageJ software, applying a threshold function to distinguish fluorescence from the background (J, L, N) and normalizing the GFP fluorescence relative to the nc82 fluorescence. Bars represent the mean of the normalized GFP fluorescence level with error bars representing the SEM. (P, W, X) Different levels of neural activity in the SOG, AL region(P), MB region (W), AL region(X) of male flies expressing *sNPF^2A^* along with the CaLexA system were observed between naïve and experienced flies. (Q-V) Quantification of GFP fluorescence in SOG (Q-R), MB (S-T), AL (U-V) between naïve and experienced *sNPF^2A^* male flies. The GFP fluorescence was processed using ImageJ software, applying a threshold function to distinguish fluorescence from the background (Q, S, V) and normalizing the GFP fluorescence relative to the nc82 fluorescence. Bars represent the mean of the normalized GFP fluorescence level with error bars representing the SEM.

The calcium levels within *sNPF^R57E07^* neurons undergo a profound shift in the brain (Fig. S6G-H), albeit with opposing trends between SOG and MB: SOG calcium decreases (Fig. 7Q-R) whereas MB calcium increases (Fig. 7S-T) in response to sexual experience. Remarkably, sexual experience does not induce a change in calcium within sNPF-positive AL neurons (Fig. 7U-V).

An intriguing observation is the substantial augmentation of calcium signals in sNPF-positive MB dAC neurons following sexual experience (Fig. 7W), implying a pivotal role for the dAC in MB-mediated gustatory input integration during interval timing-related memory processes (Lee et al. 2022; Lee et al. 2023) . Moreover, the calcium levels of sNPF-positive cells proximate to RS cells remain consistently robust under both sexually naive and experienced conditions (Fig. 7X). Collectively, these data indicate that the MB dAC and RS cells constitute pivotal neural circuits that facilitate the encoding of sexual experience-related information into memory processes through sNPF-sNPF-R peptidergic circuits (Sun et al. 2024).

### The soluble carrier transporter *Tret1l* specifically functions within sNPF neurons to regulate energy balance behavior

Despite the observation of global calcium changes in sNPF-sNPF-R circuits in response to sexual experiences, the calcium levels in sNPF neurons near antennal lobe glial cells (RS cells) did not undergo significant alterations (Fig. 7X). This suggests that metabolic shifts, rather than solely neuronal activity, may be necessary to modify the brain’s internal states within the sNPF-sNPF-R circuits. Furthermore, the consistent high calcium levels observed in sNPF neurons near RS cells indicate that these neurons rely on a significant amount of cellular energy to sustain these elevated calcium levels. Glucose metabolism in neurons is vital for their health, as it contributes to ATP production and is involved in aging and neurodegenerative diseases (Oka et al. 2021). Increased glucose transport into neurons has been shown to mitigate the toxicity of amyloid-beta (Aβ) in the fly brain (Niccoli et al. 2016), indicating that glucose metabolism is critical for maintaining brain homeostasis.

To explore the roles of sNPF-sNPF-R circuits in sensing and regulating energy balance, we focused on sugar transporters from the soluble carrier transporter (SLC) family, which control glucose metabolism between neurons and glia (Featherstone 2011; Ceder and Fredriksson 2022). Knockdown of Glucose transporter 1 (*Glut1*) in *sNPF^R57E07^* neurons significantly reduced male mating success, despite their continued courtship behavior (Fig. S6I-M). Conversely, knockdown of sugar transporter 1 (*sut1*), sugar transporter 2 (*sut2*), and Trehalose transporter 1 (*Tret1*) in sNPF neurons did not impact SMD behavior (Fig. S6N-P), indicating that these transporters are not integral to SMD behavior in sNPF-expressing neurons.

*Sut1* and *Sut2* are known to transport monosaccharides, while *Tret1* is a bidirectional trehalose transporter primarily functioning in the fat body of flies (Featherstone 2011; Volkenhoff et al. 2015; Ceder and Fredriksson 2022). The *Drosophila* genome encodes two trehalose transporters, *Tret1* and *Tret1*l, both characterized by 12 transmembrane domains (Volkenhoff et al. 2015). However, *Tret1l* lacks trehalose transporter activity (Kanamori et al. 2010; Volkenhoff et al. 2015) and its role may be less significant than *Tret1*, despite being involved in glial function (Volkenhoff et al. 2015).

Surprisingly, knockdown of *Tret1l* in *sNPF^R57E07^* neurons disrupted SMD behavior (Fig. S6Q). Knockdown of *Tret1* in *sNPF^R57E07^* neurons did not affect courtship activity after sexual experiences similar to naive and experienced wild-type males (Fig. S6R), whereas *Tret1l* knockdown reduced courtship activity (Fig. S6S). Consistent with this result, knockdown of *Tret1l* in sNPF neurons increased forward speed in males compared to naive controls (Fig. S6T-U), which is typically reduced after sexual experiences in wild-type males (Fig. S6V-W). These findings indicate that neuronal *Tret1l* finely tunes male behavior in the context of energy balance, even though its knockdown has not been reported to have significant phenotypes in previous studies (Volkenhoff et al. 2015). Glucose uptake and metabolism in neurons have been reported to be crucial for sustaining neuronal health and maintaining internal states (Besson et al. 2015; Corrales-Carvajal et al. 2016; Chatterjee and Perrimon 2021; Oka et al. 2021). Collectively, these data suggest that cellular respiration in sNPF-expressing neurons is critical for sensing and generating energy balance related to mating duration.

## DISCUSSION

In this study, we uncovered a sexually dimorphic sNPF-sNPF-R circuit in the *Drosophila* brain that regulates energy balance behavior, specifically the shorter-mating-duration (SMD) response to sexual experience. sNPF is predominantly expressed in neurons, while sNPF-R is expressed in both neurons and glial cells, particularly astrocyte-like glia (ALG) in a subset of cells outside the mushroom body (MB) termed "Rishi" cells (RS cells). Sexual experience induces global alterations in calcium signaling and synaptic plasticity within this circuit, with sNPF-R expressing neurons and glia in RS cells playing a critical role in encoding sexual experience-related information into memory processes and regulating energy balance behavior. Neuronal glucose metabolism, specifically the *Tret1l* transporter, is essential for maintaining the high calcium levels and proper function of sNPF neurons near RS cells.

The brain region that appears most comparable to RS cells is the ’proximal antennal protocerebrum (PAP)’, which is responsible for temperature coding and receives signal inputs from the antenna. The PAP is a critical component of the olfactory pathway, positioned in the anterior protocerebrum and serves as a key integrator for olfactory and higher-order sensory information (Gallio et al. 2011). Notably, the olfactory receptor neurons (ORNs) of the antenna, which highly express sNPF, provide substantial input to the PAP (Nässel et al. 2008). Considering the pivotal role of sNPF in energy balance regulation, the PAP emerges as a potential candidate for the peptidergic transmission between sNPF and its receptor, sNPF-R. However, our previous research has demonstrated that SMD behavior is not significantly affected by olfactory cues (Lee et al. 2023), indicating that the sNPF-sNPF-R circuitry from antenna to PAP may not be directly involved in SMD behavior. Therefore, it is plausible to hypothesize that the PAP region may also receive inputs from the gustatory pathway, thereby potentially modulating energy balance behavior. This intriguing aspect warrants further investigation to elucidate the complex interplay between olfaction, gustation, and energy homeostasis in the fly.

The *Drosophila* brain exhibits a sophisticated interplay between glial cells and neurons that is integral to the maintenance of energy homeostasis and behaviors (Volkenhoff et al. 2015; Endow et al. 2019; Tindell et al. 2020; Silva et al. 2022; Haynes et al. 2024). Central to this interaction is the metabolic support that glial cells offer to neurons, with a specific emphasis on IPCs (Manière et al. 2016). IPCs are equipped with specialized transporters that enable them to directly sense nutrient levels (Okamoto and Nishimura 2015), a process that can be modulated by glial cells. Astrocytes within the fly brain serve as glucose transporters, absorbing this vital energy substrate from the bloodstream and converting it into ATP through metabolic pathways such as glycolysis and the citric acid cycle (Haydon 2001). Subsequently, the generated ATP is released into the extracellular space, where it becomes accessible to neurons, including IPCs, for utilization in a variety of metabolic functions. This metabolic support is indispensable for IPCs to effectively synthesize and secrete insulin, a pivotal hormone that orchestrates energy metabolism within the brain. The synergistic relationship between glial metabolism and IPC function is thus a critical component of the neural circuitry that governs energy balance in the fly (Fig. S6H).

Insulin signaling is a fundamental regulatory mechanism in the brain, exerting its influence by modulating glucose uptake and utilization, as well as by impacting neuronal excitability and synaptic plasticity (Schwartz et al. 1992). IPCs secrete insulin into the extracellular space, where it binds to insulin receptors on target neurons. Consequently, the IPCs are essential for maintaining energy homeostasis in the fly brain. In this study, we identified a critical function of the sNPF-sNPF-R signaling pathway in the modulation of energy balance behaviors. While the expression of sNPF-R in IPCs has been previously reported (Kapan et al. 2012; Nässel et al. 2013; Suh et al. 2015) and our study provides further confirmation (Fig. 3Q), the precise mechanisms by which the sNPF-sNPF-R circuitry regulates energy homeostasis through IPCs remain largely elusive.

The IPCs in adult *Drosophila* are capable of directly detecting elevated sugar concentrations in the hemolymph, triggering the secretion of *Drosophila* insulin-like peptides (dILPs) (Nässel and Zandawala 2020). The secretion of dILPs from IPCs is modulated by a variety of neuronal inputs. Notably, sNPF, secreted by sugar-sensing CN-neurons has been shown to stimulate the secretion of dILP2 from IPCs (Oh et al. 2019). These findings implicate sNPF receptors in IPCs as crucial for glucose sensing and insulin secretion. Our investigation revealed that sNPF-R expression in IPCs is also crucial for modulating energy balance behaviors (Fig. 3). However, it should be noted that the sNPF receptor-expressing IPCs do not correspond to the fru-positive, male-specific cell population (Fig. 6A).

In conclusion, our findings reveal a novel and intricate neuron-glia sNPF-sNPF-R network that acts as an accumulator circuit within the *Drosophila* brain for processing interval timing behaviors (Gibbon 1977; Kacelnik and Brunner 2002; Rijn et al. 2014).

This network integrates pacemaker information from sNPF-expressing clock neurons in the RS cells and transmits energy-related internal states to the MB memory circuits (Sun et al. 2024). This integration is crucial for generating interval timing behaviors, particularly in male sexual behavior, highlighting the dynamic interplay between metabolism, neural circuits, and behavior in regulating energy balance related to mating duration.

## ACKNOWLEDGMENTS

We thank Drs. Yuh Nung Jan and Lily Yeh Jan (UCSF, USA) for helpful comments and support on this paper. We are also very appreciative to the colleagues who supplied us with several fly strains: Dr. Wei Zhang (Tsinghua University), Fang Guo (Zhejiang University), and Dr. Yufeng Pan (Southeast University) and Drs. Young-Joon Kim and Sung-Eun Yoon (Korea Drosophila Resource Center, KDRC). This research was supported a University of Ottawa Startup grant 602496 to WJK, Startup funds from HIT Center for Life Science to WJK, a University of Ottawa Interdisciplinary Research Group Funding Opportunity (IRGFO stream 1 and 2) grants 148101 and 148747 to WJK, a Natural Sciences and Engineering Research Council of Canada (NSERC) Discovery grant (reference: 211406) to WJK, a University of Ottawa Brain and Mind Research Institute/Center for Neural Dynamics Open call project grant 150950 to WJK, a Mitacs Globalink Research Internship Program grant 17268 to WJK. This research was also supported by the Brain Pool Program of the National Research Foundation in Korea grant ZYM5041911 to WJK, Burroughs Wellcome Fund Collaborative Research Travel Grants (reference: 1017486) to WJK and a NVIDIA Academic Hardware Grant Program to WJK. The funders had no role in study design, data collection and analysis, decision to publish, or preparation of the manuscript.

## DECLARATION OF GENERATIVE AI AND AI-ASSISTED TECHNOLOGIES IN THE WRITING PROCESS

During the creation of this work, the author(s) utilized QuillBot to rephrase English sentences, verify English grammar, and detect plagiarism, as none of the authors of this paper are native English speakers. After using this tool/service, the author(s) reviewed and edited the content as needed and take(s) full responsibility for the content of the publication.

## SUPPLEMENTAL INFORMATION AUTHOR CONTRIBUTIONS

**Conceptualization:** Woo Jae Kim.

**Data curation:** Xiaoli Zhang, Hongyu Miao, Dayeon Kang, Dongyu Sun, and Woo Jae Kim.

**Formal analysis:** Xiaoli Zhang, Hongyu Miao, Dayeon Kang, Dongyu Sun, and Woo Jae Kim.

**Funding acquisition:** Woo Jae Kim.

**Investigation:** Woo Jae Kim.

**Methodology:** Woo Jae Kim.

**Project administration:** Woo Jae Kim.

**Resources:** Woo Jae Kim.

**Supervision:** Woo Jae Kim.

**Validation:** Xiaoli Zhang, Woo Jae Kim. **Visualization:** Xiaoli Zhang, Woo Jae Kim. **Writing – original draft:** Woo Jae Kim.

**Writing – review & editing:** Xiaoli Zhang, Woo Jae Kim.

**STAR** IZ **METHODS**

## RESOURCE AVAILABILITY

### Lead contact

Further information and requests for resources and reagents should be directed to and will be fulfilled by the lead contact, Woo Jae Kim.

## EXPERIMENTAL MODEL AND STUDY PARTICIPANT DETAILS

### Drosophila

*Drosophila melanogaster* were cultured under standard laboratory conditions at 25℃. Samples were prepared as described in the Methods Details. All fly strains are listed in the key resources table.

## METHOD DETAILS

### Fly Stocks and Husbandry

*Drosophila melanogaster* were raised on cornmeal-yeast medium at similar densities to yield adults with similar body sizes. Flies were kept in 12 h light: 12 h dark cycles (LD) at 25[ (ZT 0 is the beginning of the light phase, ZT12 beginning of the dark phase) except for some experimental manipulation (experiments with the flies carrying tub-GAL80^ts^). Wild-type flies were Canton-S. To reduce the variation from genetic background, all flies were backcrossed for at least 3 generations to CS strain. All mutants and transgenic lines used here have been described previously. Following lines used in this study, *Df*(1)*Exel6234* (#7708)*, sNPF^[c00448]^* (#85000)*, Pdf ^[01]^* (#26654)*, lexAop-CD8GFP; UAS-mLexA-VP16-NFAT, lexAop-rCD2-GFP* (#66542),

*lexAop-nSyb-spGFP1-10, UAS-CD4-spGFP11* (#64315)*, NPFR^[c01896]^* (#10747),

*MB247-GAL4* (#50742)*, ey[ok107]-GAL4* (#854)*, dilp2-GAL4* (#37516)*, repo-GAL4*

(#7415)*, sNPF-GAL4^57E07^* (#46382)*, sNPF-GAL4^64H09^* (#46547)*, elav^c155^* (#458),

*UAS-traF* (#4590)*, fru-GAL4* (#66696)*, UAS>stop>TNT* (#67690)*, fruFlp* (#66870),

*dsx-GAL4* (#63434)*, UAS>stop>KCNJ2* (#67686)*, sNPF-R^[MI00427]^* (#30996),

*UAS-CD4tdGFP* (#35839)*, UAS-RedStinger* (#8546)*, sNPF-RNAi^JF01906^* (#25867),

*sNPF-R-RNAi^JF02657^*(#27507)*, UAS-Denmark, UAS-syt.eGFP* (#33065)*, UAS-KCNJ2*

(#6596)*, UAS-mCD8RFP, lexAop-mCD8GFP* (#32229)*, UAS-TNT* (#28838),

*UAS-myrGFP.QUAS-mtdTomato-3Xha; trans-Tango* (#77124)*, ALG-GAL4* (#45914),

*CG-GAL4* (#39944)*, SPG-GAL4* (#50472)*, PNG-GAL4* (#40436)*, EGT-GAL4*

(#39908)*, EGN-GAL4* (#39157)*, pdf-GAL4* (#84683)*, cry-GAL4* (#24774),

*ClK4.1M-GAL4* (#36316)*, VGlut-GAL4* (#41275)*, Gad1-GAL4* (#51630),

*UAS-NaChBac* (#9467)*, ChAT-RNAi^JF01877^* (#25856)*, Gad1-RNAi^JF02916^* (#28079),

*VGlut-RNAi^JF02689^*(#27538)*, Gad1-RNAiJF02916* (#28079)*, UAS-DenMark* (#33062) were obtained from the Bloomington *Drosophila* Stock Center at Indiana University. The following lines, *UAS>stop>Kir2.1/CyO; Dsx-Flp* (#1186)*, UAS>stop>mCD8GFP; Dsx-Flp* (#1172)*, UAS-sNPF-R* (#10066)*, UAS>stop>Shi^ts^, dsx-FLP* (#1386)*, w-; UAS>stop>DSCAM17.1-GFP; fruFlp* (#1122)*, UAS-sNPF* (#10064)*, MB247-GAL80* (#2001)*, Cry-GAL80* (#2680)*, UAS>stop>TNT ^inactive^*

(#1124)*, UAS>stop>Kir2.1/CyO; Dsx-Flp* (#1186)*, UAS>stop>mCD8GFP; Dsx-Flp*

(#1172)*, UAS-sNPF-R* (#10066)*, UAS>stop>Shi^ts^, dsx-FLP* (#1386)*, w-;*

*UAS>stop>DSCAM17.1-GFP; fruFlp* (#1122)*, UAS>stop>TNT ^inactive^* (#1124),

*UAS-sNPF* (#10064)*, MB247-GAL80* (#2001)*, Cry-GAL80* (#2680) were obtained from Korea Drosophila Resource Center (KDRC). The following line, *Cha-GAL4* was obtained from Qidong Fungene Biotechnology in China. The *UAS>stop>mCD8GFP; fruFlp* and *UAS>stop>nSyb-GFP; fruFlp* were a gift from Dr. Barry Dickson (HHMI Janelia Research Campus, USA). The *UAS>stop>ricin* was a gift from Dr. David J. Anderson (Howard Hughes Medical Institute).

### Mating duration assay

The mating duration assay in this study has been reported(Kim et al. 2012; Kim et al. 2013; Lee et al. 2023). To enhance the efficiency of the mating duration assay, we utilized the *Df(1)Exel6234* (DF here after) genetic modified fly line in this study, which harbors a deletion of a specific genomic region that includes the sex peptide receptor (SPR)(Parks et al. 2004; Yapici et al. 2008). Previous studies have demonstrated that virgin females of this line exhibit increased receptivity to males(Yapici et al. 2008). We conducted a comparative analysis between the virgin females of this line and the CS virgin females and found that both groups induced SMD. Consequently, we have elected to employ virgin females from this modified line in all subsequent studies. For naïve males, 40 males from the same strain were placed into a vial with food for 5 days. For experienced males, 40 males from the same strain were placed into a vial with food for 4 days then 80 DF virgin females were introduced into vials for last 1 day before assay. 40 DF virgin females were collected from bottles and placed into a vial for 5 days. These females provide both sexually experienced partners and mating partners for mating duration assays. At the fifth day after eclosion, males of the appropriate strain and DF virgin females were mildly anaesthetized by CO_2_. After placing a single female in to the mating chamber, we inserted a transparent film then placed a single male to the other side of the film in each chamber. After allowing for 1 h of recovery in the mating chamber in 25[ incubator, we removed the transparent film and recorded the mating activities. Only those males that succeeded to mate within 1 h were included for analyses. Initiation and completion of copulation were recorded with an accuracy of 10 sec, and total mating duration was calculated for each couple. Genetic controls with *GAL4/+* or *UAS/+* lines were omitted from supplementary figures, as prior data confirm their consistent exhibition of normal LMD and SMD behaviors (Kim *et al*, 2013, 2012; Huang *et al*, 2024; Lee *et al*, 2023; Zhang *et al*, 2024). Hence, genetic controls for SMD behavior was incorporated exclusively when assessing novel fly strains that had not previously been examined. In essence, internal controls were predominantly employed in the experiments, as SMD behavior exhibit enhanced statistical significance when internally controlled. In the context of SMD experiments, the naïve condition and sexually experienced males act as mutual internal controls for one another. A statistically significant divergence between naïve and experienced males indicates that the experimental procedure does not alter SMD. Conversely, the absence of a statistically significant difference suggests that the manipulation does impact SMD. Hence, we incorporated supplementary genetic control experiments solely if they deemed indispensable for testing. All assays were performed from noon to 4 PM. We conducted blinded studies for every test.

### Immunostaining

After 5 days of eclosion, the Drosophila brain was taken from adult flies and fixed in 4% formaldehyde at room temperature for 30 minutes. The sample was than washed three times (5 minutes each) in 1% PBT and then blocked in 5% normal goat serum for 30 minutes. Subsequently, the sample was incubated overnight at 4℃ with primary antibodies in 1% PBT, followed by the addition of fluorophore-conjugated secondary antibodies for one hour at room temperature. Finally, the brain was mounted on plates with an antifade mounting solution (Solarbio) for imaging. purposes.

### Quantitative analysis of fluorescence intensity

To quantify the calcium level and synaptic intensity in microscopic images, we introduced ImageJ software(Wiki.). We initially employed ImageJ’s ‘Measure’ feature to calculate average pixel intensity across the entire image or in user-specified sections, and the ‘Plot Profile’ feature to create intensity profiles across areas. To maximize precision, we converted color images to grayscale before analysis.

Thresholding methods were also utilized to produce binary images that accurately outlined areas of interest, with pixel intensities of 255 (white) assigned to regions of interest and 0 (black) to the background. Intensity values from the binary image were then transferred to the corresponding locations in the original grayscale image to obtain precise intensity measurements for each object. The ‘Display Results’ feature provided comprehensive data for each object, including average intensity, size, and other relevant statistics. To normalize for fluorescence differences between ROIs, GFP fluorescence for CaLexA was normalized to nc82. All specimens were imaged under identical conditions.

### Colocalization analysis

To perform colocalization analysis of multi-color fluorescence microscopy images in this study, we employed ImageJ software(Daly. 2020). Initially, merge the individual channels of the fluorescence image to create a composite view. Following this, access the ‘Image’ menu, select ‘Type’, and then choose ‘RGB Color’ to ensure the image is displayed in the correct color format. Next, navigate to ‘Image’, then ‘Adjust’, and select ‘Color Threshold’. Adjust the threshold settings to isolate the yellowish pixels indicative of colocalization, and confirm the selection by clicking the ‘Select’ button. Subsequently, proceed to ‘Analyze’, followed by ‘Measure’. This action will open a new window displaying the measurement data for the selected area. Record the ‘Area’ value provided in the measurement window, as this corresponds to the area of colocalization between the two fluorophores. To calculate the percentage of the colocalized area relative to the total area of the fluorophore of interest (e.g., GFP or RFP), readjust the threshold settings to encompass the entire fluorophore area and click ‘Select’ again. Afterward, repeat the measurement process by going to ‘Analyze’ and then ‘Measure’. This will yield the total area value for the respective fluorophore. Finally, to determine the colocalization percentage, divide the area value of the colocalized region by the total area value of the GFP or RFP region. This calculation provides the colocalization efficiency within the specified region. Additionally, we displayed images of fluorescence and overlapping areas in the male fly brain and/or VNC, processed with ImageJ software using a threshold function to differentiate fluorescence from background. All specimens were imaged under identical conditions.

### Particle analysis

To quantitatively measure particle intensity of cell number and synaptic puncta in microscopic images, we applied ImageJ software(Wiki.). Initially, the image is converted to grayscale to reduce complexity and enhance contrast. Subsequently, thresholding techniques are employed to binarize the image, distinguishing particles from the background. This binarization can be achieved through automated thresholding algorithms or manual adjustment to optimize the segmentation.

Following binarization, the “Analyze Particles” function is utilized to enumerate and measure the particles based on user-defined criteria, such as size constraints and shape descriptors (e.g., area, perimeter, circularity). The results of these measurements are then available for review in the “Results” window, and the particles can be labeled within the image for visual identification. All specimens were imaged under identical conditions.

### Single-nucleus RNA-sequencing analyses

snRNAseq dataset analyzed in this paper is published(Li et al. 2022) and available at the Nextflow pipelines (VSN, https://github.com/vib-singlecell-nf), the availability of raw and processed datasets for users to explore, and the development of a crowd-annotation platform with voting, comments, and references through SCope (https://flycellatlas.org/scope), linked to an online analysis platform in ASAP (https://asap.epfl.ch/fca). Single-cell RNA sequencing (scRNA-seq) data from the *Drosophila melanogaster* were obtained from the Fly Cell Atlas website (https://doi.org/10.1126/science.abk2432). Oenocytes gene expression analysis employed UMI (Unique Molecular Identifier) data extracted from the 10x VSN oenocyte (Stringent) loom and h5ad file, encompassing a total of 506,660 cells. The Seurat (v4.2.2) package (https://doi.org/10.1016/j.cell.2021.04.048) was utilized for data analysis. Violin plots were generated using the “Vlnplot” function, the cell types are split by FCA.

### Statistical Tests

Statistical analysis of mating duration assay was described previously(Kim et al. 2012; Kim et al. 2013; Lee et al. 2023). More than 50 males (naïve, experienced and single) were used for mating duration assay. Our experience suggests that the relative mating duration differences between naïve and experienced condition and singly reared are always consistent; however, both absolute values and the magnitude of the difference in each strain can vary. So, we always include internal controls for each treatment as suggested by previous studies(Bretman et al. 2011). Therefore, statistical comparisons were made between groups that were naïvely reared, sexually experienced and singly reared within each experiment. As mating duration of males showed normal distribution (Kolmogorov-Smirnov tests, p > 0.05), we used two-sided Student’s t tests. The mean ± standard error (s.e.m) (*\*\*\*\* = p < 0.0001, *** = p < 0.001, ** = p < 0.01, * = p < 0.05*). All analysis was done in GraphPad (Prism). Individual tests and significance are detailed in figure legends.

Besides traditional *t*-test for statistical analysis, we added estimation statistics for all MD assays and two group comparing graphs. In short, ‘estimation statistics’ is a simple framework that—while avoiding the pitfalls of significance testing—uses familiar statistical concepts: means, mean differences, and error bars. More importantly, it focuses on the effect size of one’s experiment/intervention, as opposed to significance testing(Claridge-Chang and Assam 2016). In comparison to typical NHST plots, estimation graphics have the following five significant advantages such as (1) avoid false dichotomy, (2) display all observed values (3) visualize estimate precision (4) show mean difference distribution. And most importantly (5) by focusing attention on an effect size, the difference diagram encourages quantitative reasoning about the system under study(Ho et al. 2019). Thus, we conducted a reanalysis of all of our two group data sets using both standard *t* tests and estimate statistics. In 2019, the Society for Neuroscience journal eNeuro instituted a policy recommending the use of estimation graphics as the preferred method for data presentation(Bernard 2021).

### Locomotion Assay

To image fly locomotion, a custom LED array was positioned beneath the stage to serve as backlighting. A 2[mm transparent acrylic board sheet was then placed atop the LED array to ensure homogeneous illumination. Fly behavior was recorded using a camera with a resolution of 1920×1080 pixels at 30 frames per second (1080p30). Behavioral arenas were custom-built from opaque transparent acrylic board sheets.

The chambers measured 30[mm in diameter and 2[mm in height. Group flies were aspirated into each chamber and positioned on the stage. A 30-minute observation period was conducted to ensure the absence of gross motor defects before initiating video acquisition.

### Video Acquisition and Tracking

Videos were captured using a camera with a resolution of 1920×1080 pixels at 30 frames per second (1080p30). Image segmentation was performed using custom software called Fly Trajectory Dynamics Tracking (FlyTrDT), developed in Python. FlyTrDT identifies and quantifies fruit flies as elliptical pixels. The main feature extracted from the videos was the average forward speed of the fly group within each chamber. For the average forward speed of group flies, FlyTrDT records the movement speed of each indoor fruit fly and analyzes the average speed of 10 fruit flies per second.

## SUPPLEMENTAL FIGURE TITLES AND LEGENDS

Figure S1. **sNPF-to-sNPF-R circuits are composed of functional neuron-glia network. (**A) SMD assay of *sNPF^c00448^*/+. (B) Expression patterns of *sNPF^2.5^* male brain. (D-E) SMD assay of SMD assays for *sNPF-RNAi/+* (D), *sNPF-R-RNAi/+* (E). (H-K) SMD assays were performed for *GAL4* mediated overexpression of *sNPF via sNPF^R57E07^*(H) and *sNPF-R^R64H09^* (I) and for overexpression of *sNPF-R via sNPF^R57E07^*(J) and *sNPF-R^R64H09^* (K). (L-M) SMD assays for *UAS-sNPF /+* (L), *UAS-sNPF-R /+* (M).

Figure S2. **A distinct subset of cells outside of mushroom body that express the sNPF-R is integral to the regulation of SMD behavior.** (A-C) SMD assays for *GAL4^MB247^/+* (A)*, GAL4^ok107^/+* (B)*, GAL4^ok107^*, *elav^c155^*, *GAL4^MB247^/+* (C). (D) Quantification of cell number. Bars represent the mean *GAL4^ok107^* (black column), *GAL4^MB247^* (white column), the combination *GAL4^ok107^*, *GAL80^MB247^* (red column) and *dilp2-GAL4* (gray column) cell number fluorescence level with error bars representing SEM. (E) SMD assays of *dilp2-GAL4*/+. (F-I) Brain of male flies expressing expressing *sNPF^R57E07^* (F) and *sNPF^R57E07^*, *GAL80^MB247^* (G), *sNPF-R^R64H09^* (H) and *sNPF-R^R64H09^, GAL80^MB247^* (I) together with *UAS-Redstinger.* (J) Quantification of cell number of male fly brain expressing *sNPF^R57E07^* and *sNPF^R57E07^*, *GAL80^MB247^* driver together with *UAS-Redstinger*. (K) Expression pattern of *sNPF-R^R64H09^*, SPG^R54C07^ and ALG^R86E01^ in Virtual Fly Brain (VFB) and the colored box indicate their co-location.

Figure S3. T**he astrocyte-like glial expression of sNPF-R is essential for the manifestation of SMD behavior.** (A-F) SMD assays for *ALG-GAL4*/+ (A)*, CG-GAL4*/+ (B)*, EGN-GAL4*/+ (C)*, EGT-GAL4*/+ (D)*, PNG-GAL4*/+ (E), *SPG-GAL4*/+ (F). (G-L) SMD assays were conducted to activate *ALG-GAL4* (G)*, CG-GAL4* (H)*, EGN-GAL4* (I)*, EGT-GAL4* (J)*, PNG-GAL4* (K), *SPG-GAL4* (L) sodium channel crossed with *UAS-NaChBac*. (M-N) SMD assays for *UAS-NaChBac/+ and deathstar-GAL4.* (O-P) Locomotion assay comparing the knockdown of *ALG-GAL4* in *mCherry-RNAi* with the knockdown of *ALG-GAL4* in *deathstar-RNAi* in males (O) and females (P). (Q) Diagram of connectivity between *sNPF-sNPF-R and deathstar*.

Figure S4. **The sexually dimorphic nature of sNPF-sNPF-R neural circuits constitutes a pivotal regulatory mechanism for SMD behavior.** (A) Quantification of *sNPF-R^R64H09^* total area of brain between males (blue column) and feminized males (yellow column), displaying fluorescence levels with error bars representing SEM. (B) Brain of male and female flies expressing *sNPF^R57E07^* together with *UAS-CD4tdGFP, UAS-Redstinger.* (C-F) Quantification of cell number (D), total area (E) and average size (F) between male and female. The flies are described as (B). The RFP fluorescence was processed using ImageJ software, applying a threshold function to distinguish fluorescence from the background (C). (G) Brain of male flies expressing *sNPF ^R57E07^* with *UAS-FRT.stop-mCD8GFP, fru^FLP^* were immunostained with anti-GFP (green), and nc82 (magenta) antibodies. Scale bars represent 100 μm. Boxes indicate the magnified regions of interest presented in the middle panels. (H) Quantification of cell number between CB and OL. The flies are described as (G).

Figure S5. **Neuronal anatomy and structural features of RS cells.** (A-B) SMD assays for *sNPF^R57E07^* cross with *UAS-FRT.stop-ricin, fru^FLP^* (A), *UAS-FRT.stop-TNT^in^, fru^FLP^* (B). (C-D) SMD assays for *sNPF-R^R64H09^* cross with *UAS-FRT.stop-ricin, fru^FLP^* (A), *UAS-FRT.stop-TNT^in^, fru^FLP^* (B). (E) SMD assay of *sNPF-R^R64H09^* cross with *UAS-traF*. (F) SMD assays of *fru-GAL4* mediated knockdown of *sNPF-R via sNPF-R-RNAi*. (G-H) Brain of male flies expressing *GAL4^MB247^* and *GAL4^ok107^*. (I) Brain of male flies expressing *GAL4^ok107^*, *GAL80^MB247^* with *UAS-syteGFP, UAS-denmark* were immunostained with anti-GFP (green), anti-DsRed (red) and nc82 (blue) antibodies. Scale bars represent 100 μm. (J-K) Expression patterns of RS region for *sNPF^R57E07^*(J), *sNPF-R^R64H09^*(K) in the Virtual Fly Brain (VFB). (L) Co-localization analysis of *sNPF^R57E07^* and *sNPF-R^R64H09^* in Virtual Fly Brain (VFB). The yellow area represents overlap by *sNPF-R^R64H09^* (green) and *sNPF^R57E07^* (red). (N-O) Expression patterns of *beat-VII^R94F06^* (M), *sNPF-R^R64H09^* (N) in the Virtual Fly Brain (VFB). (P) Co-localization analysis of *beat-VII^R94F06^* and *sNPF-R^R64H09^* in Virtual Fly Brain (VFB). The yellow area represents overlap by (green) and *sNPF-R^R64H09^* (red). (Q-S) Boxes (Q-S) indicate the magnified regions of interest presented in the middle. The flies are described as (N-P). The yellow area represents overlap by *beat-VII^R94F06^* (green) and *sNPF-R^R64H09^* (red) in RS region.

Figure S6. **Sexual experience induces global alterations in calcium signaling throughout sNPF-sNPF-R brain circuits.** (A-D) Different levels of neural activity in the CB and AL regions of male flies expressing *GAL4^ok107^* along with the CaLexA system were observed between naïve and experienced flies. (E-F) Different levels of neural activity in the brain of male flies expressing *sNPF-R^R64H09^* along with the CaLexA system were observed between naïve and experienced flies. (G-H) Different levels of neural activity in the brain of male flies expressing *sNPF^R57E07^* along with the CaLexA system were observed between naïve and experienced flies. (I) SMD assay for *sNPF^R57E07^* cross with *Glut1-RNAi*. (J-K) Comparison of mating success ratio between naïve and experienced *sNPF^R57E07^* crossed with control-RNAi (J) and *sNPF^R57E07^* crossed with *Glut1-RNAi* (K). (L-M) Comparison of courtship index between *sNPF^R57E07^* crossed with control-RNAi and *sNPF^R57E07^* crossed with *Glut1-RNAi*. (N-O) SMD assay for *sNPF^R57E07^* cross with *Sut1*-RNAi(N) and *Sut2*-RNAi(O). (P-Q) SMD assay for *sNPF^R57E07^* cross with *Tret1*-RNAi(P) and *Tret1*l-RNAi(Q). (R-S) Comparison of courtship index *sNPF^R57E07^* cross with *Tret1*-RNAi(R) and *Tret1*l-RNAi(S). (T-U) Locomotion assay of *sNPF^R57E07^* cross with *Tret1*-RNAi. (V-W) Locomotion assay of *sNPF^R57E07^* cross with control-RNAi.

Table 1**. Screen of 14 GAL4s.**

