## Supplementary figures and images for "Male-specific sNPF peptidergic circuits control energy balance for mating duration through neuron-glia interactions"

### Supplemental figure 1

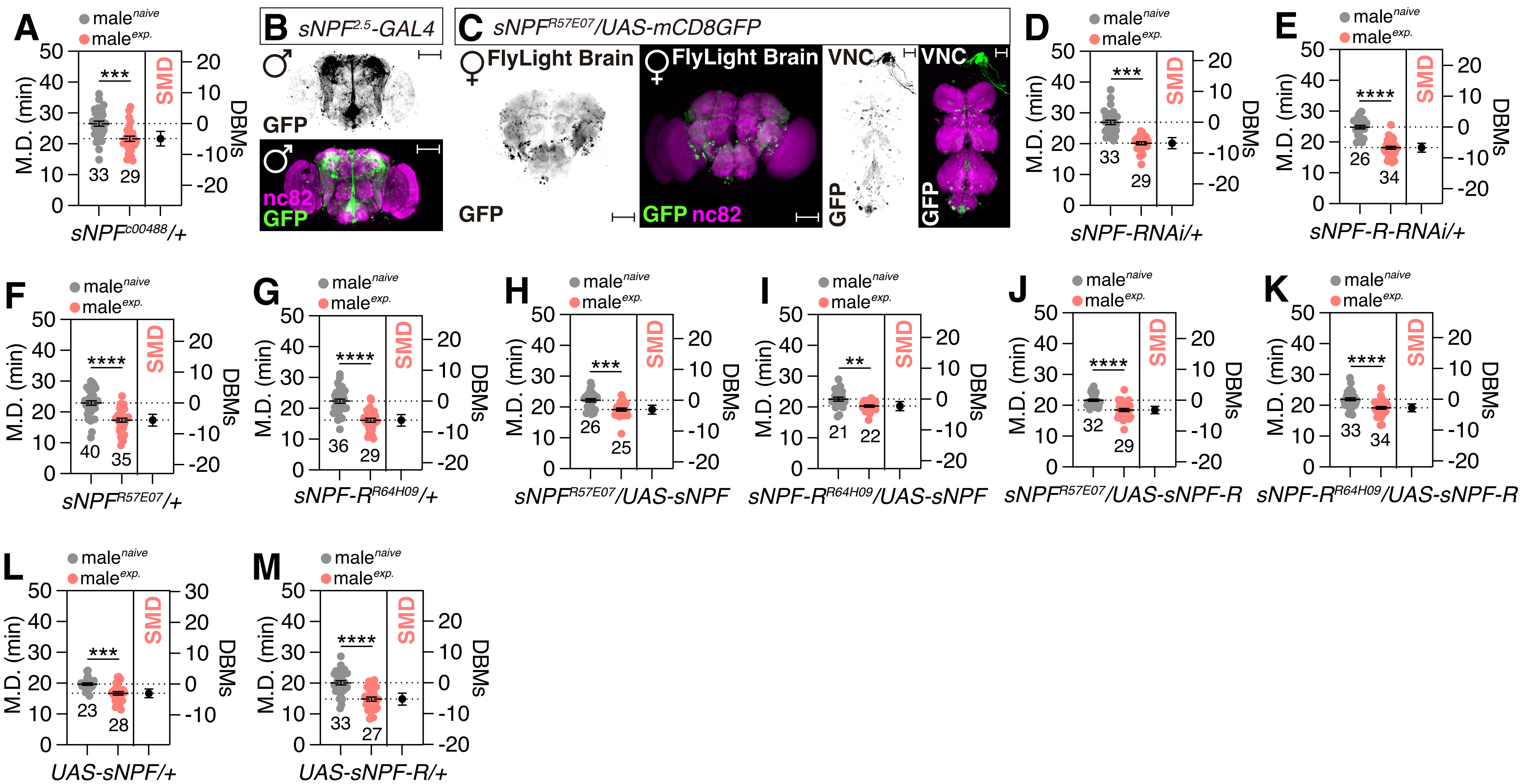

### Supplemental figure 2

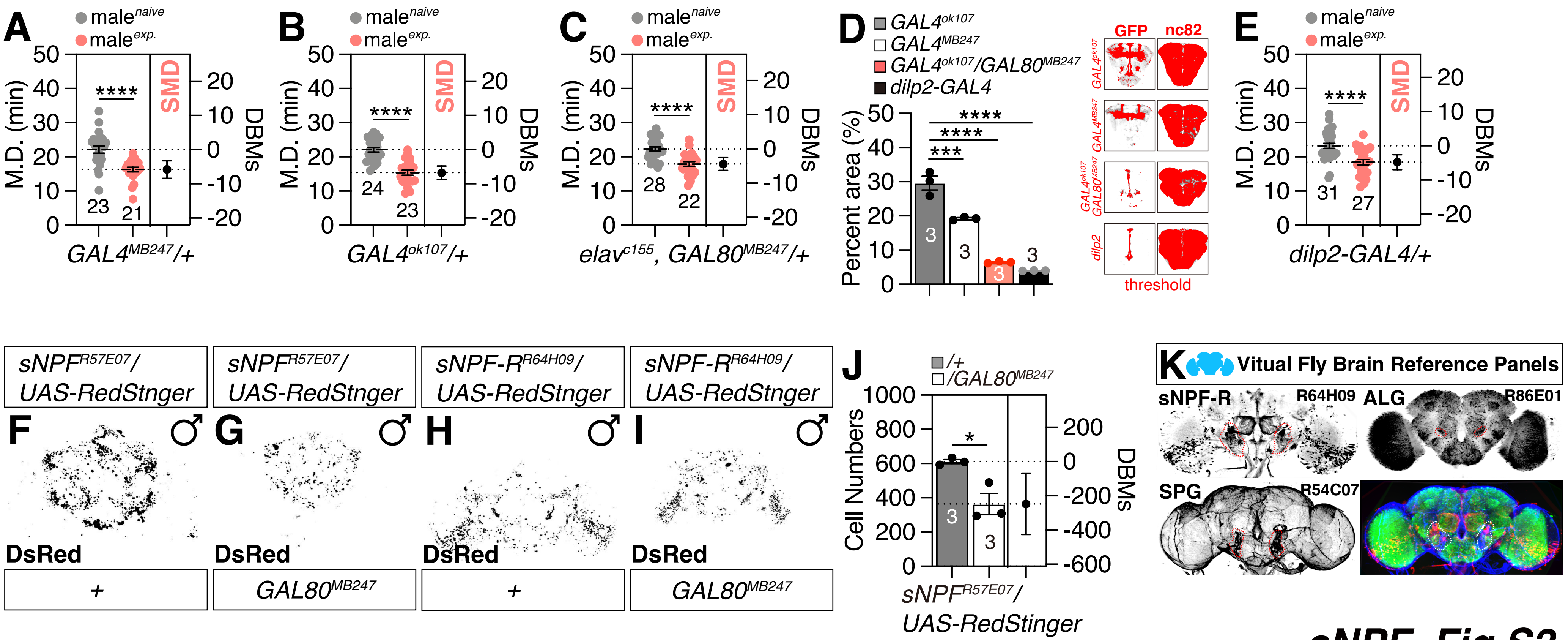

### Supplemental figure 3

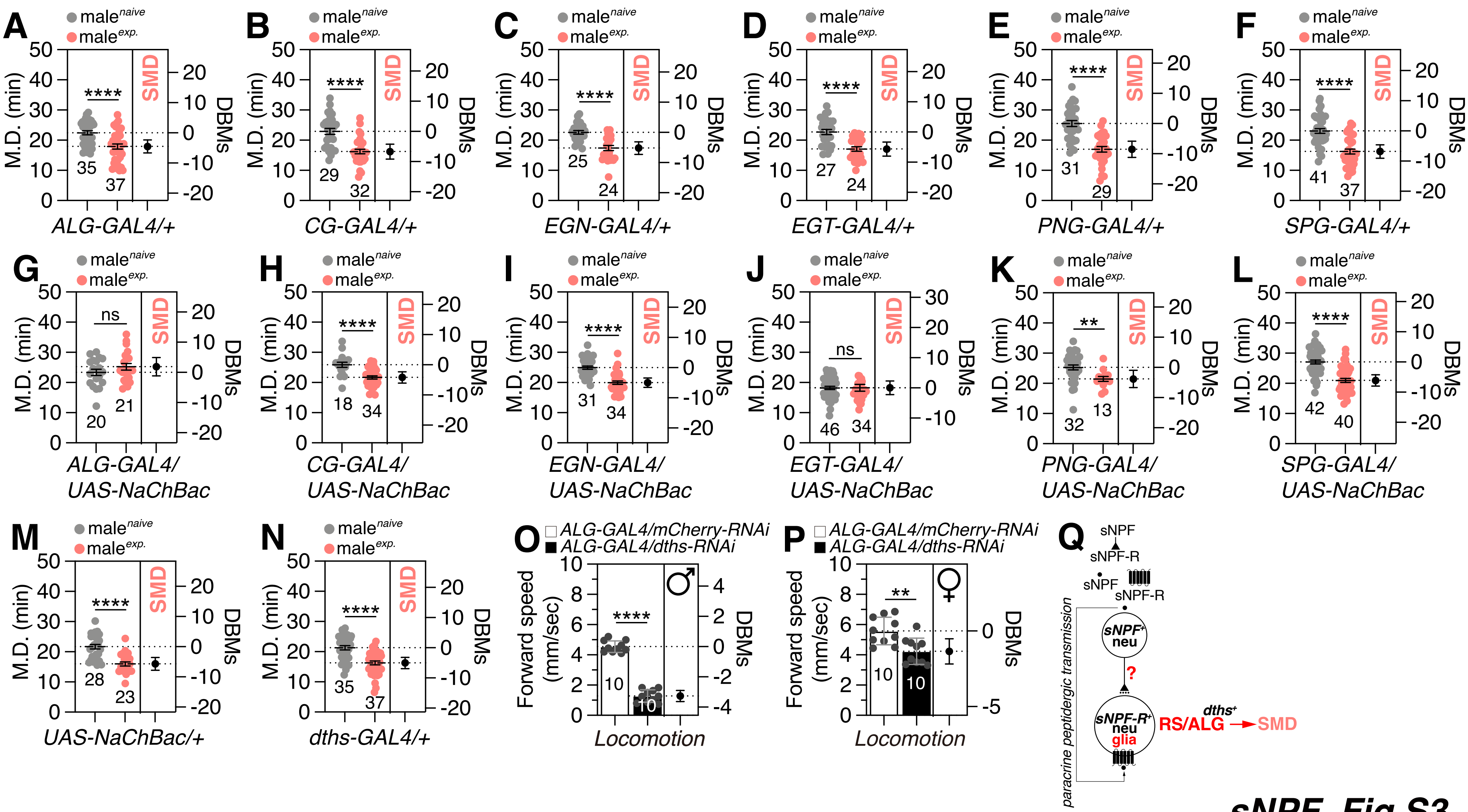

### Supplemental figure 4

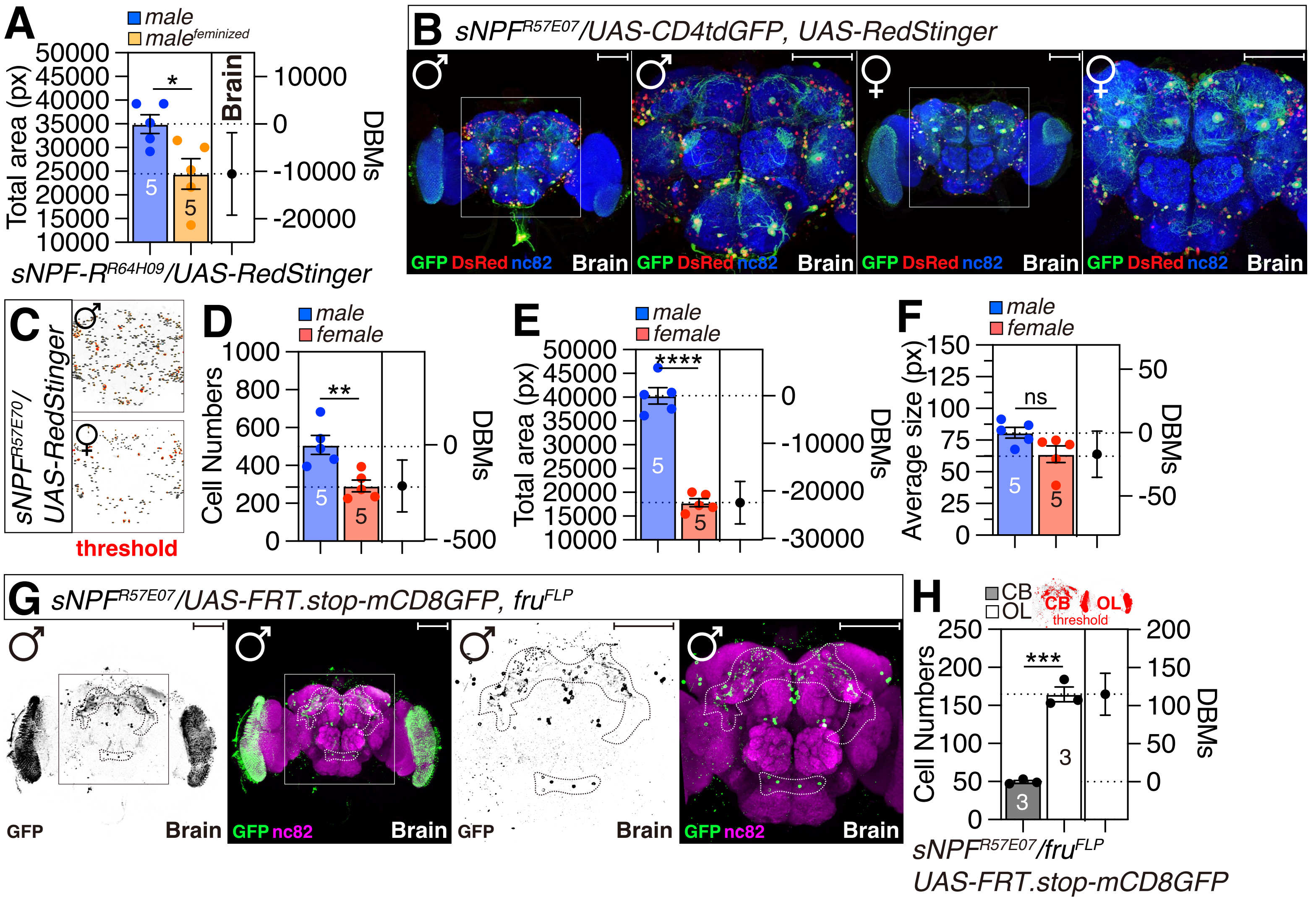

### Supplemental figure 5

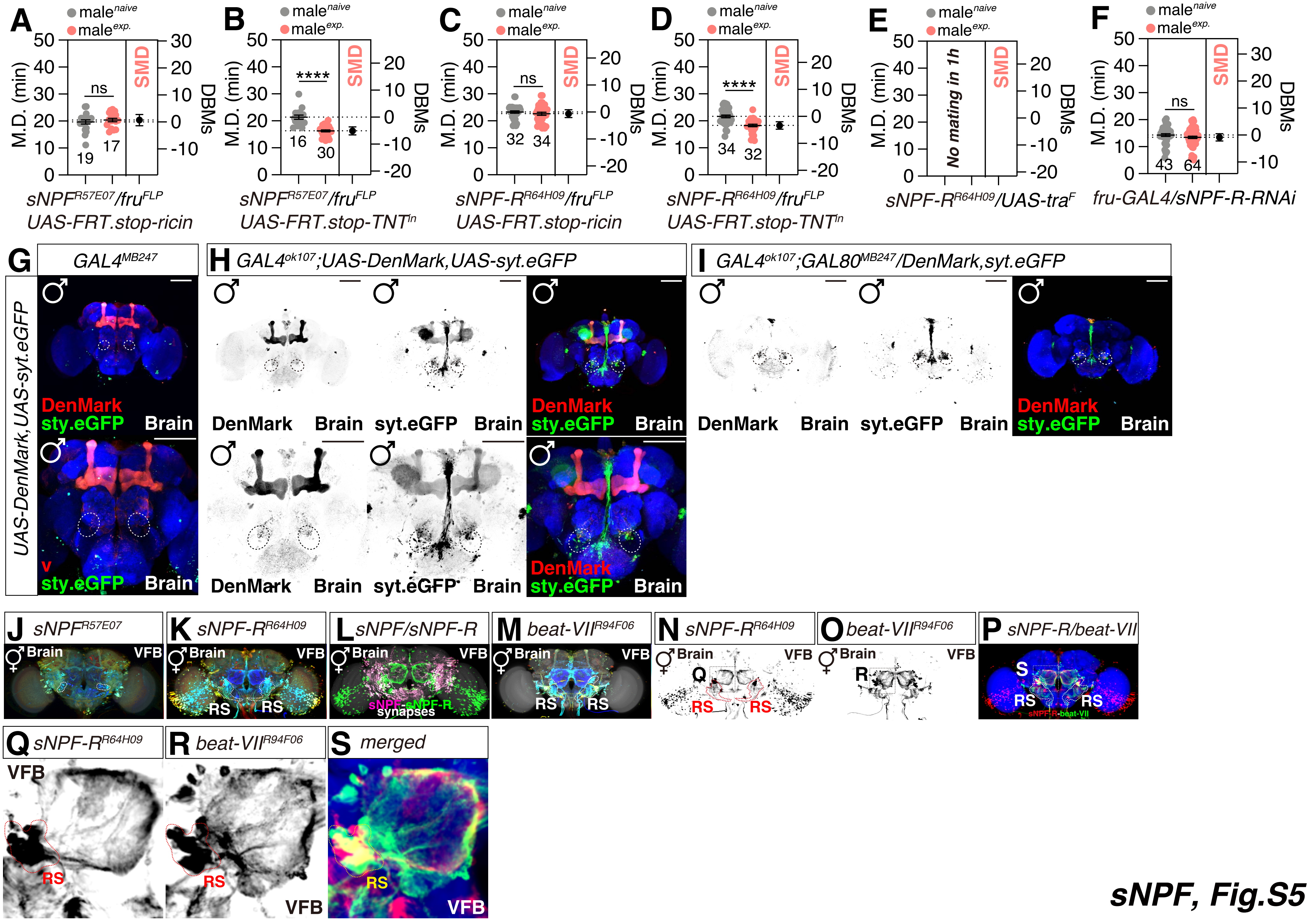

### Supplemental figure 6

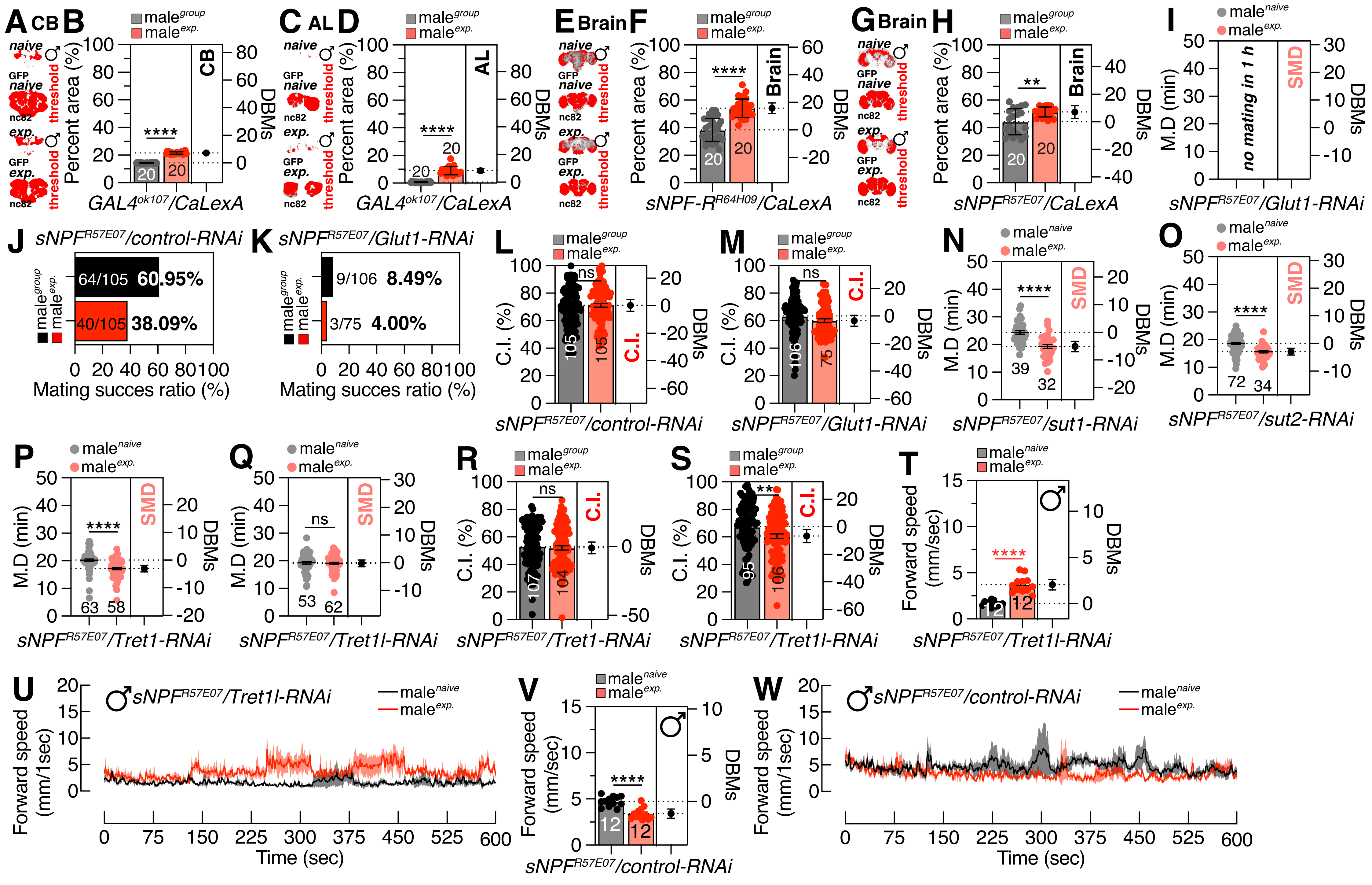
