## Supplemental table 1 for "Male-specific sNPF peptidergic circuits control energy balance for mating duration through neuron-glia interactions"

**Table S1** Screening GAL4s mediated knockdown of sNPF-R via sNPF-R-RNAi.

| **GAL4** | **Genotype** | **SMD** | **T-test** | **Gene expression** |
| --- | --- | --- | --- | --- |
| *R19H12* | *GAL4^R19H12^/*  *UAS-sNPF-R-RNAi* | ns | 0.4822 | 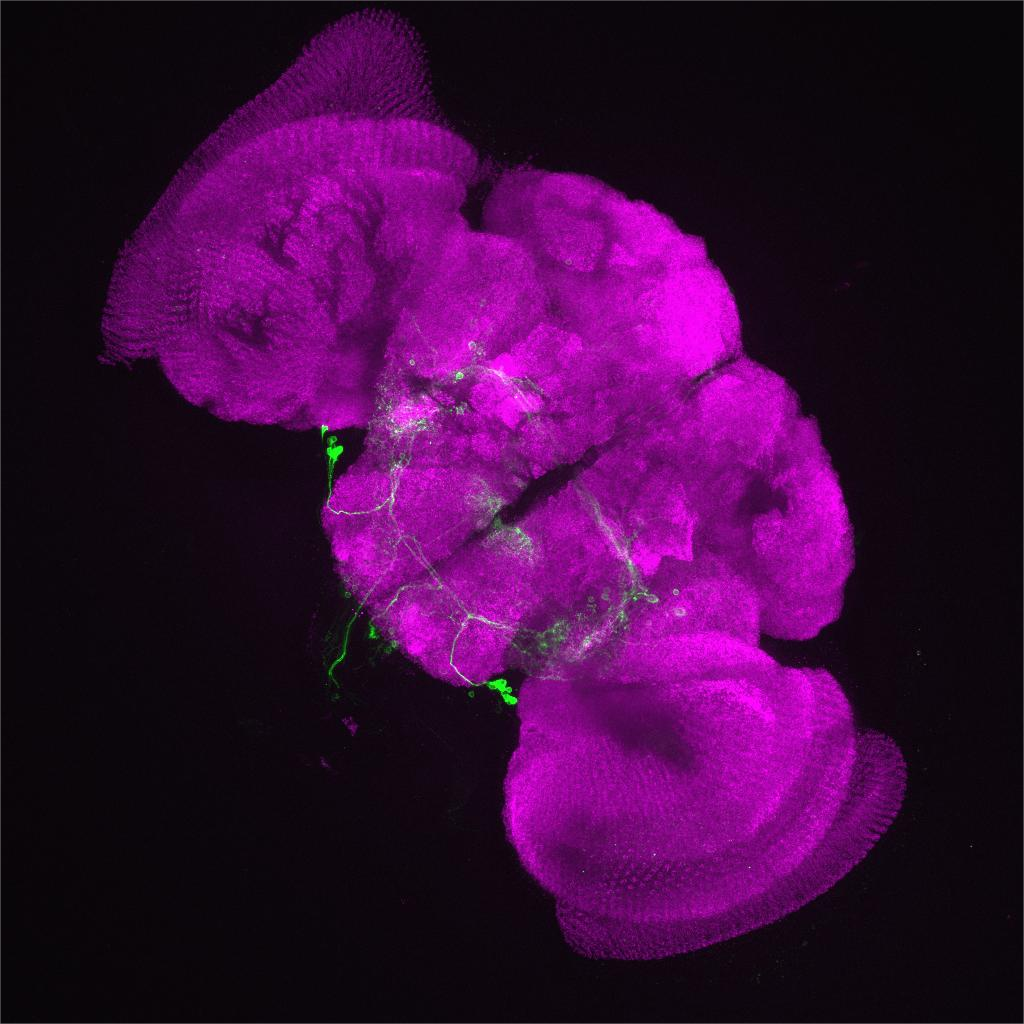 |
| R20B12 | *GAL4^R20B12^/*  *UAS-sNPF-R-RNAi* | ** | 0.0054 | 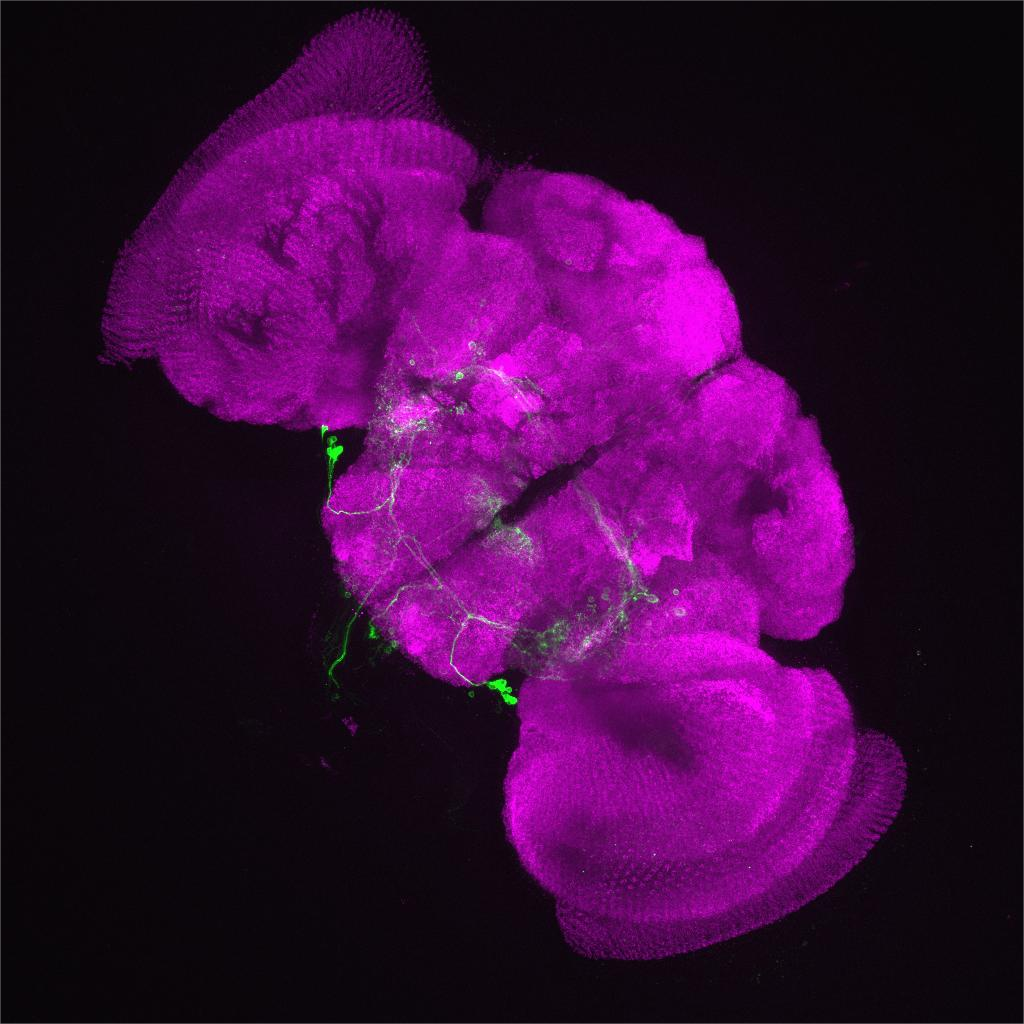 |
| R20D06 | *GAL4^R20D06^/*  *UAS-sNPF-R-RNAi* | ns | 0.5795 | 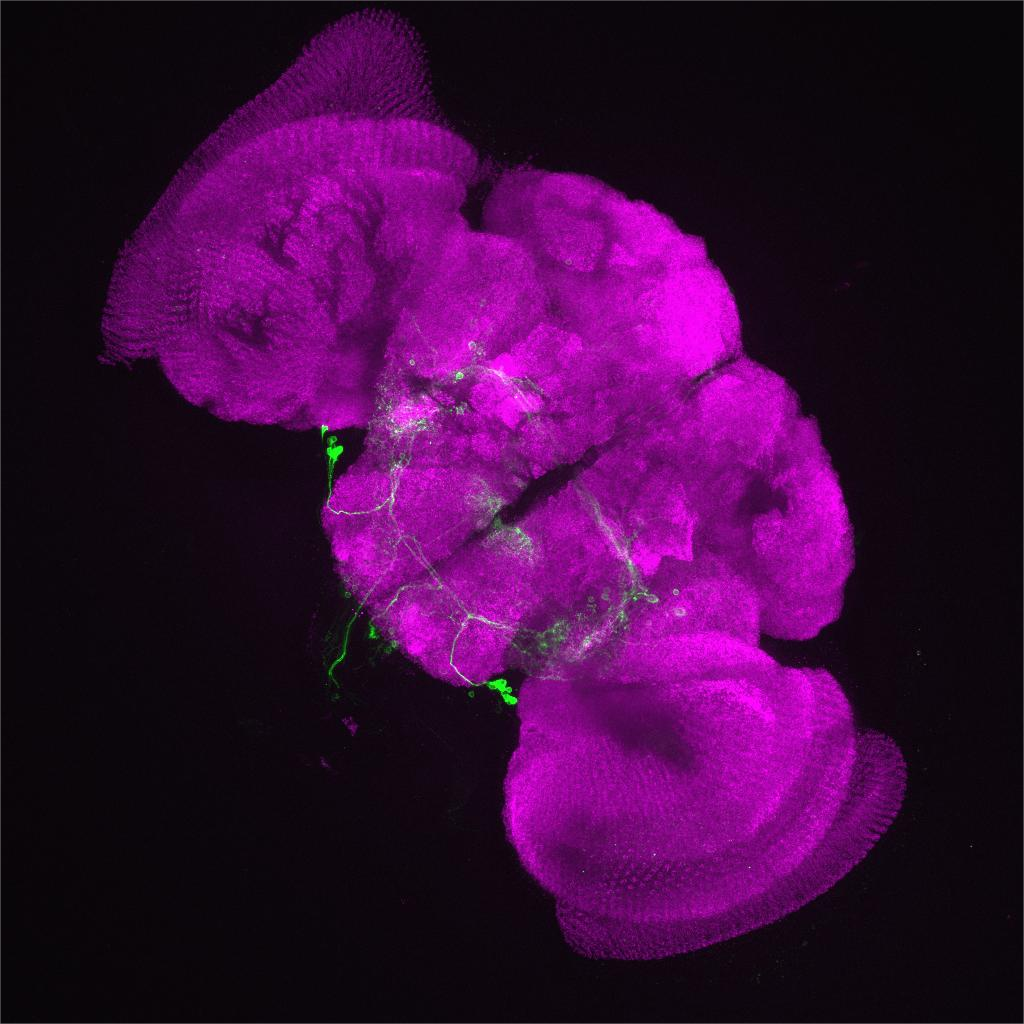 |
| R20E08 | *GAL4^R20E08^/*  *UAS-sNPF-R-RNAi* | ns | 0.2970 | 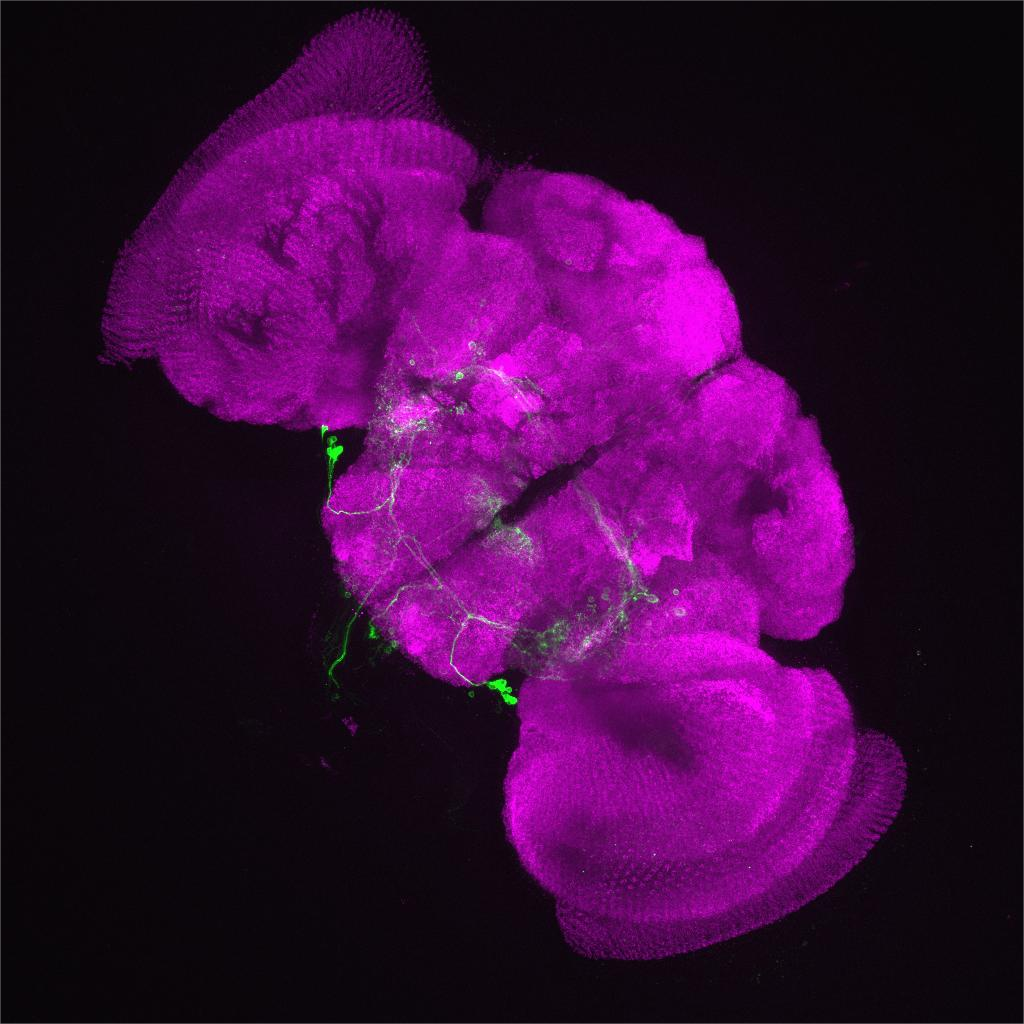 |
| R20F11 | *GAL4^R20F11^/*  *UAS-sNPF-R-RNAi* | **** | <0.0001 | 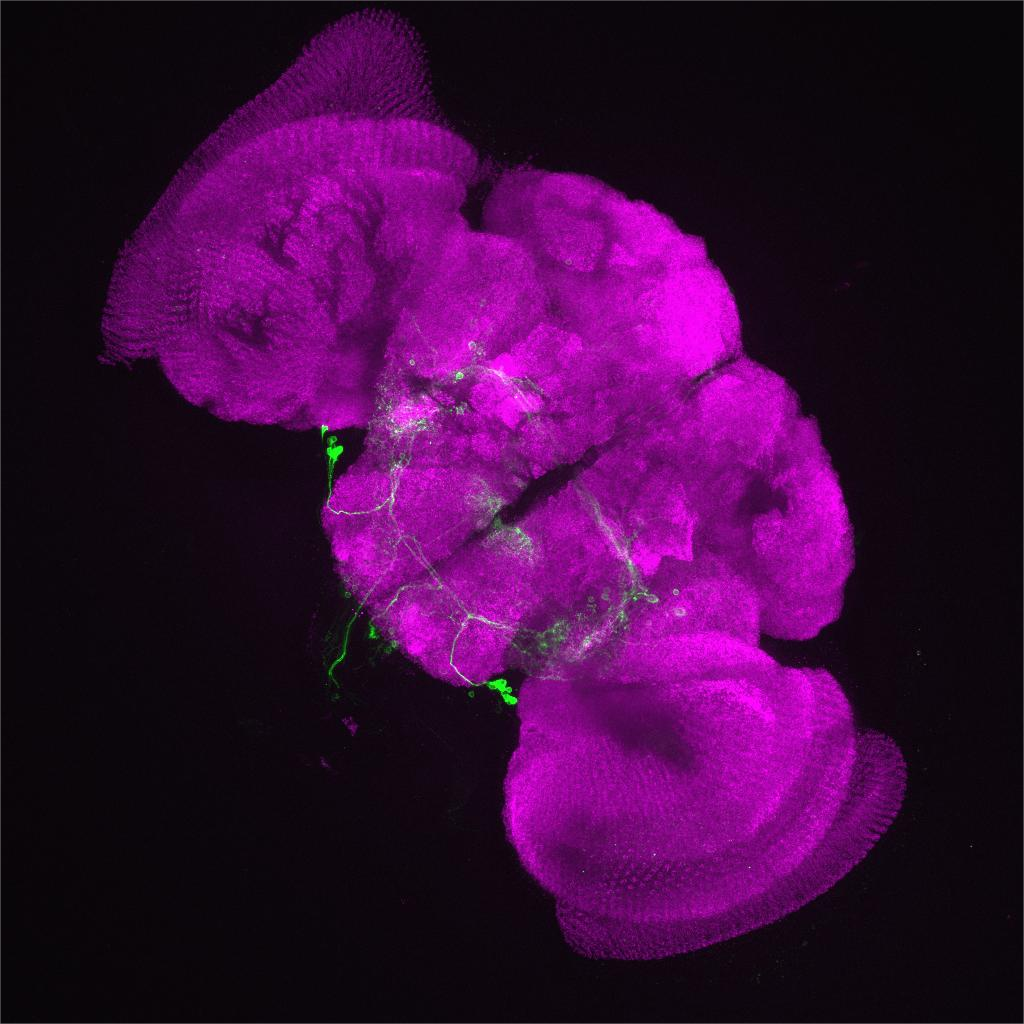 |
| R21A12 | *GAL4^R21A12^/*  *UAS-sNPF-R-RNAi* | **** | <0.0001 | 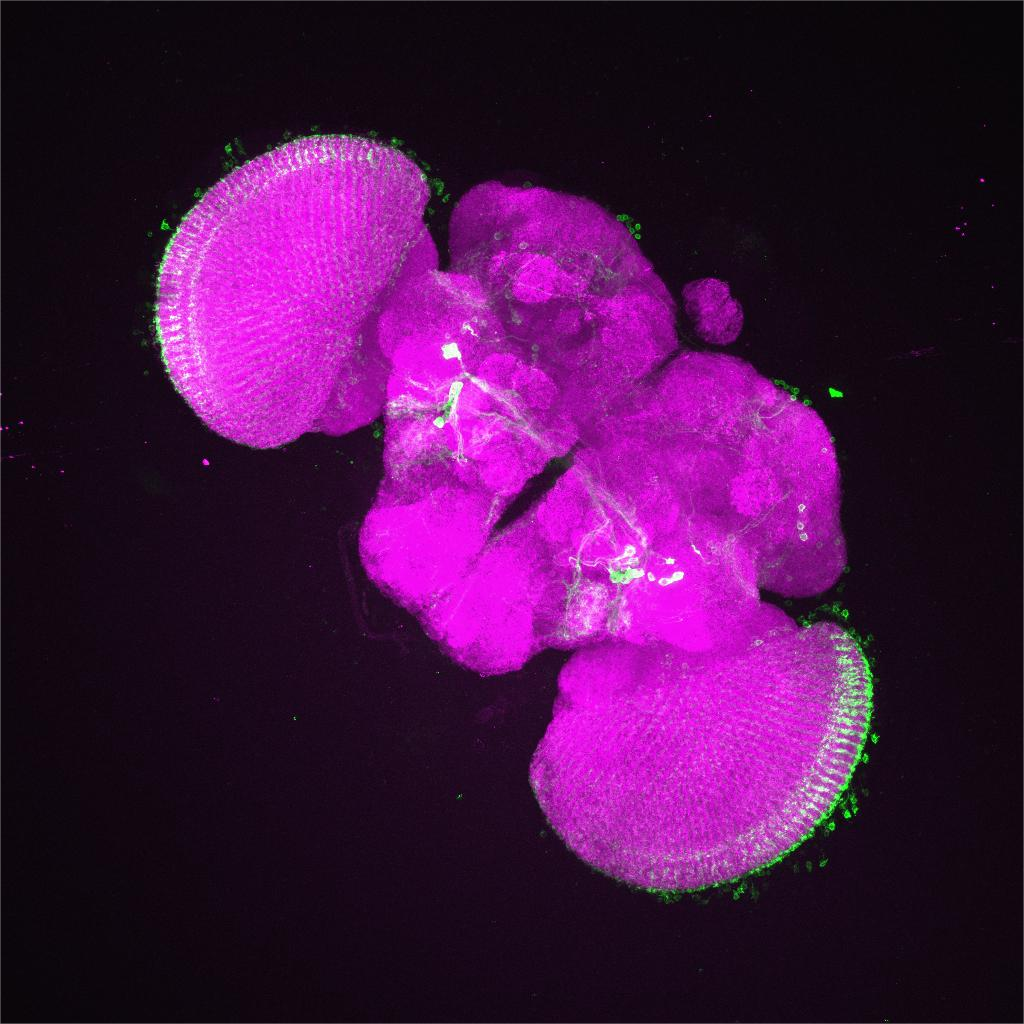 |
| R21B01 | *GAL4^R21B01^/*  *UAS-sNPF-R-RNAi* | *** | 0.00020 | 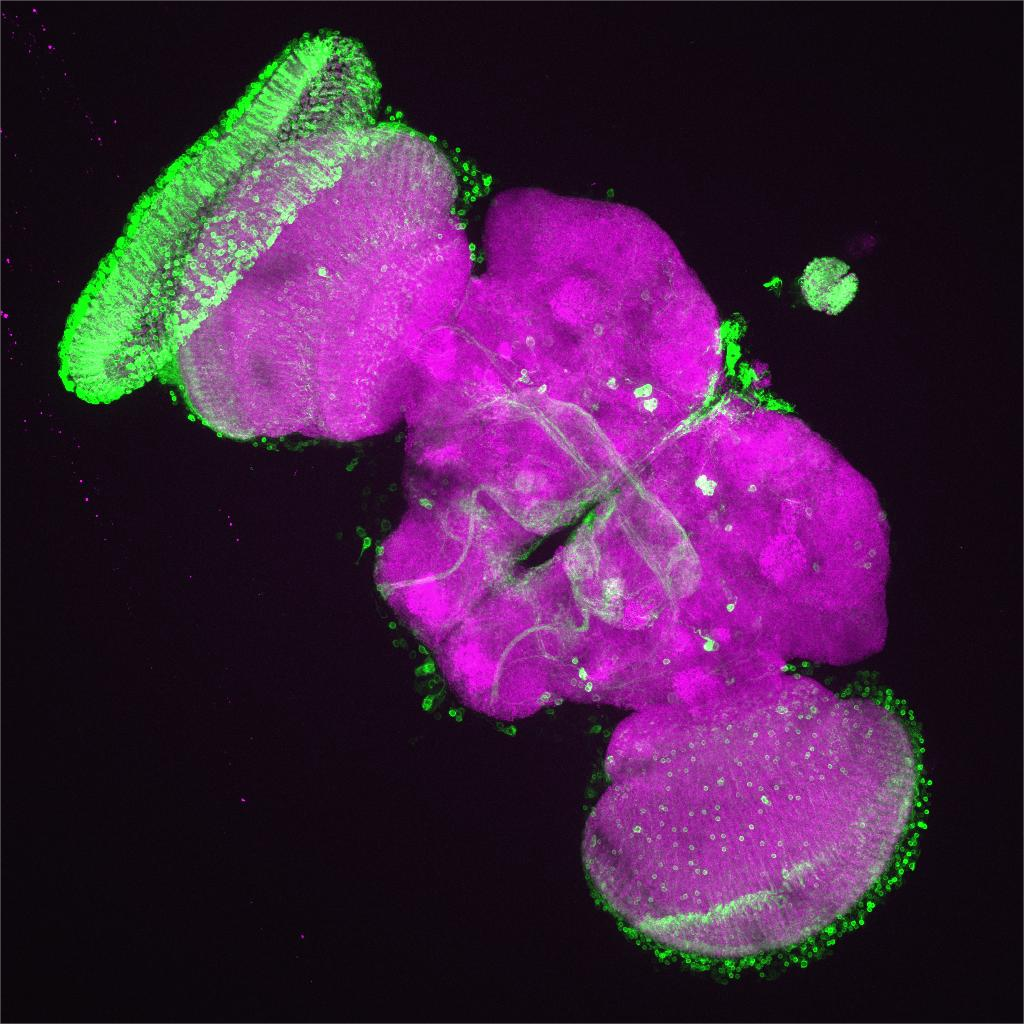 |
| R21C05 | *GAL4^R21C05^/*  *UAS-sNPF-R-RNAi* | ns | 0.9203 | 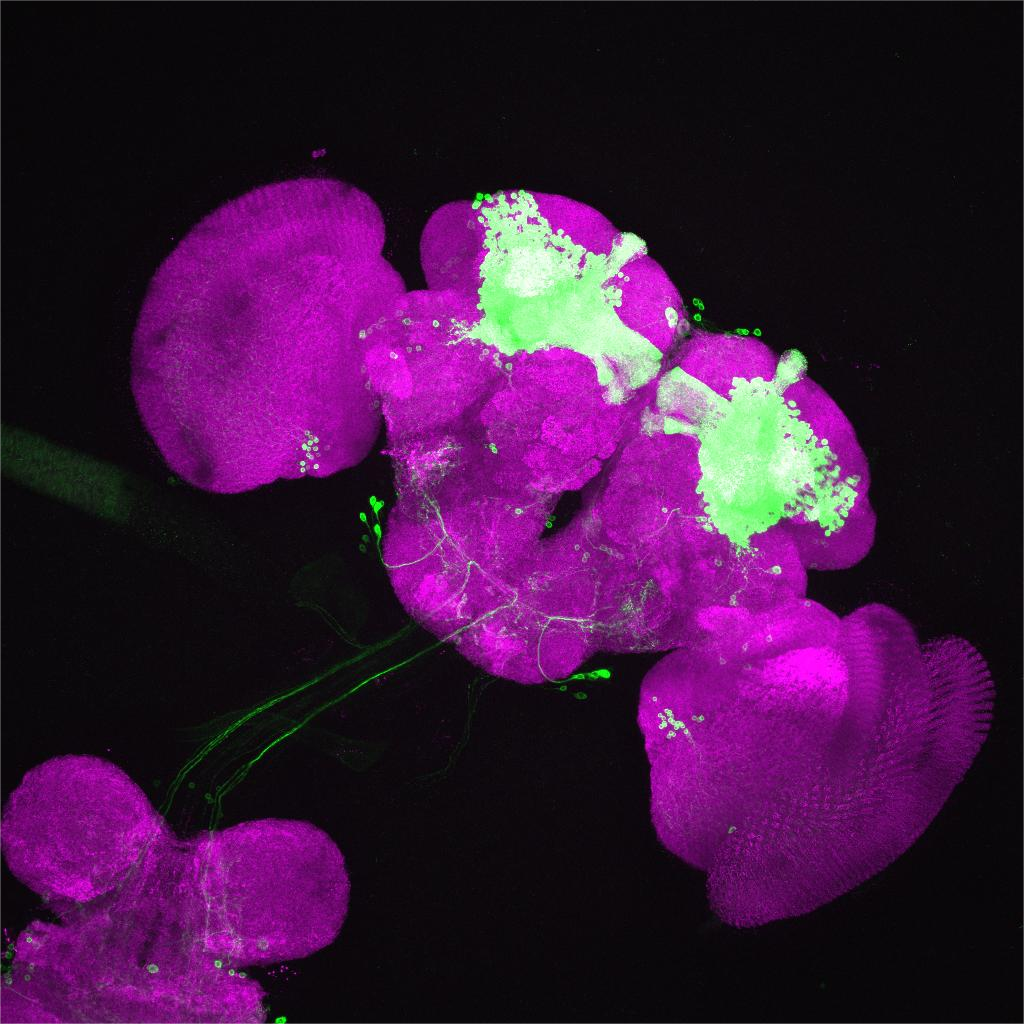 |
| R64A06 | *GAL4^R64A06^/*  *UAS-sNPF-R-RNAi* | **** | <0.0001 | 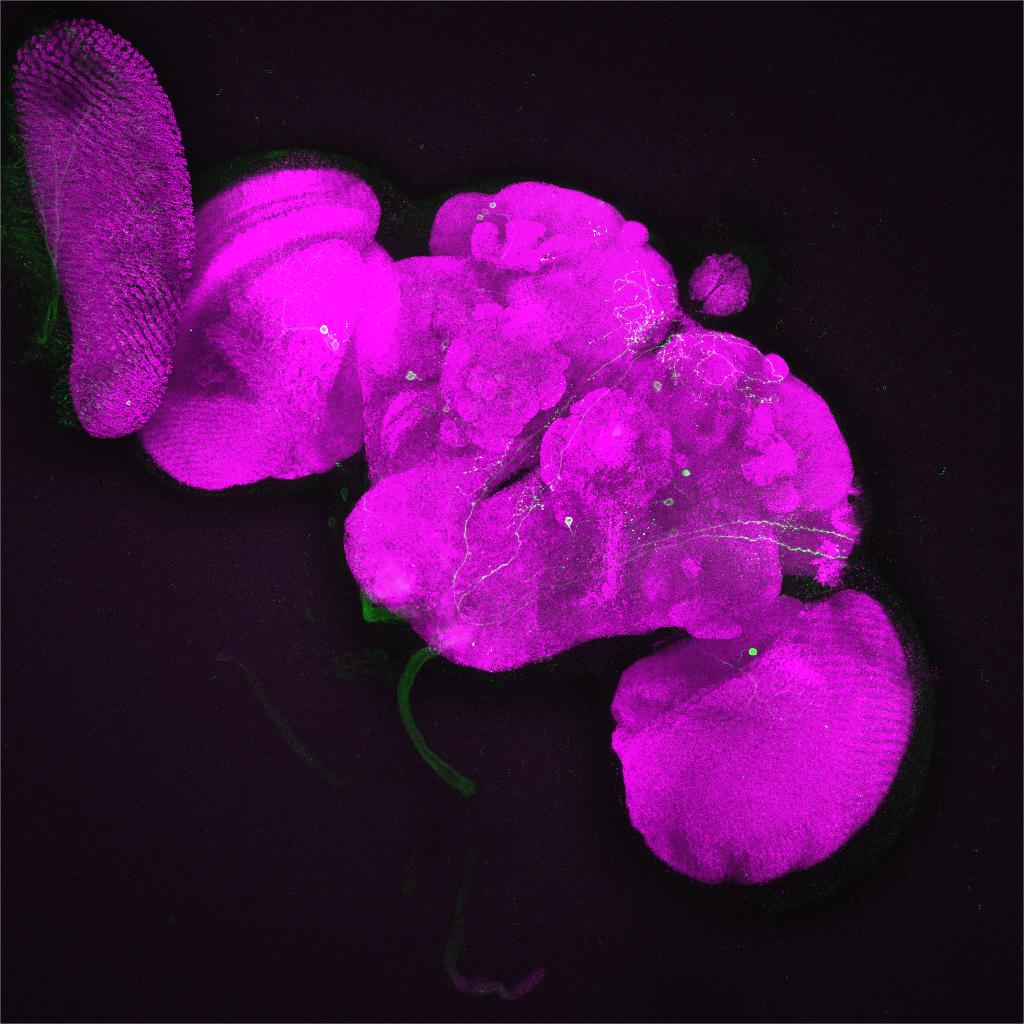 |
| R64B11 | *GAL4*^R64B11^*/*  *UAS-sNPF-R-RNAi* | ns | 0.1998 | 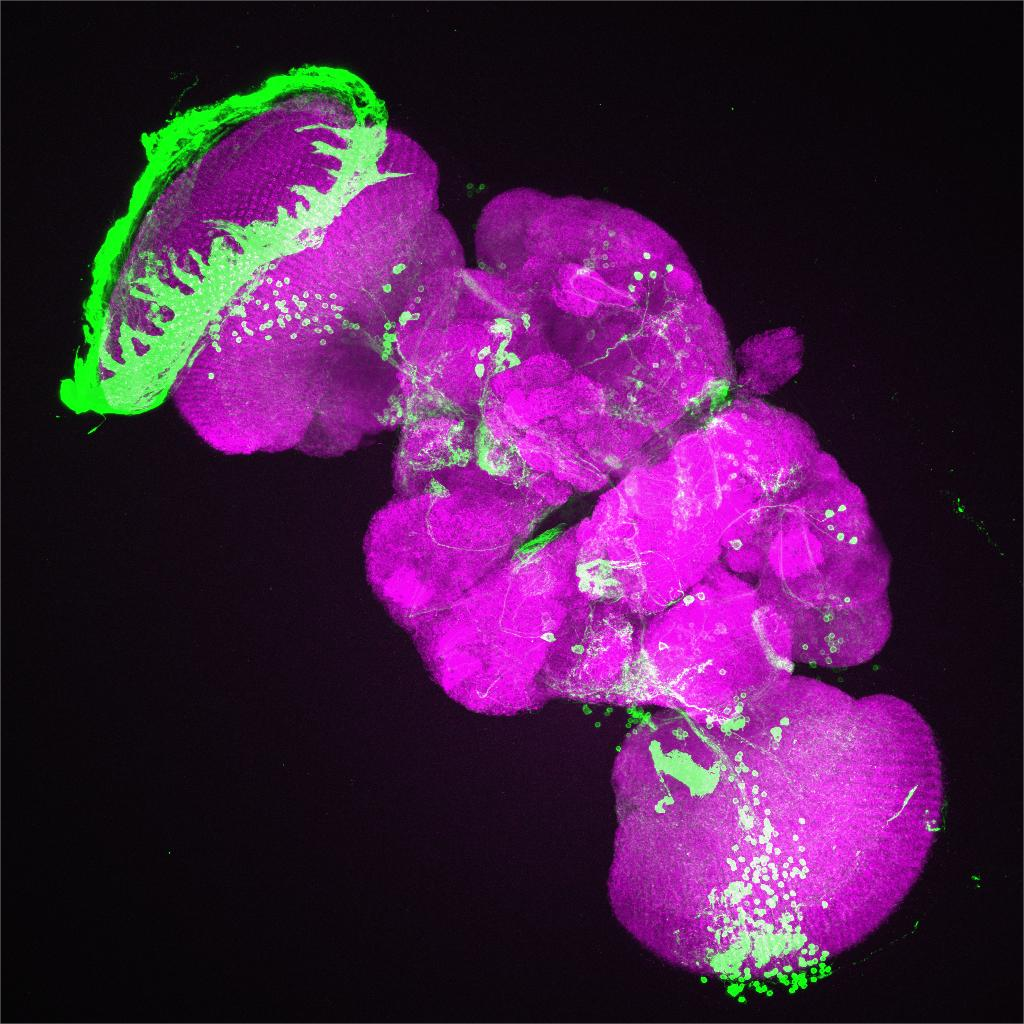 |
| R64D05 | *GAL4*^R64D05^*/*  *UAS-sNPF-R-RNAi* | ** | 0.0072 | 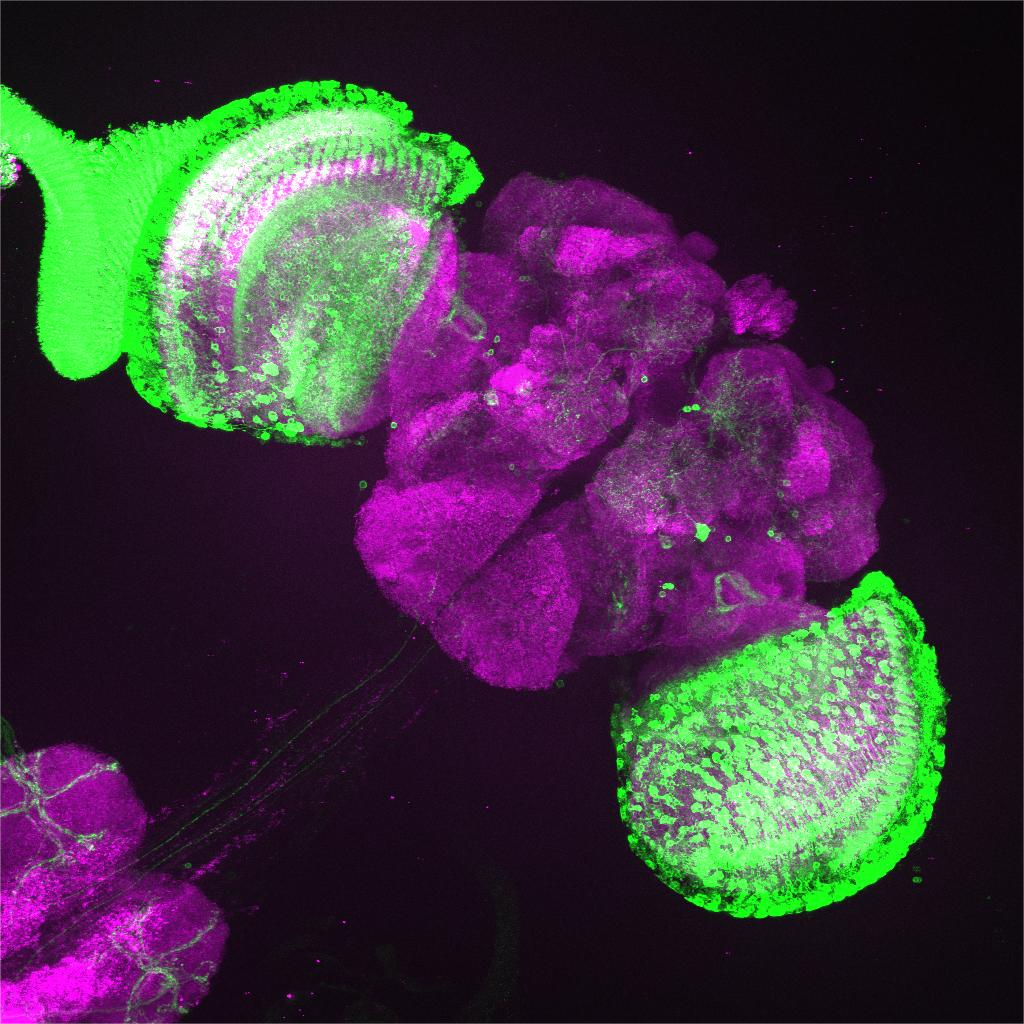 |
| R64F07 | *GAL4*^R64F07^*/*  *UAS-sNPF-R-RNAi* | ns | 0.5253 | 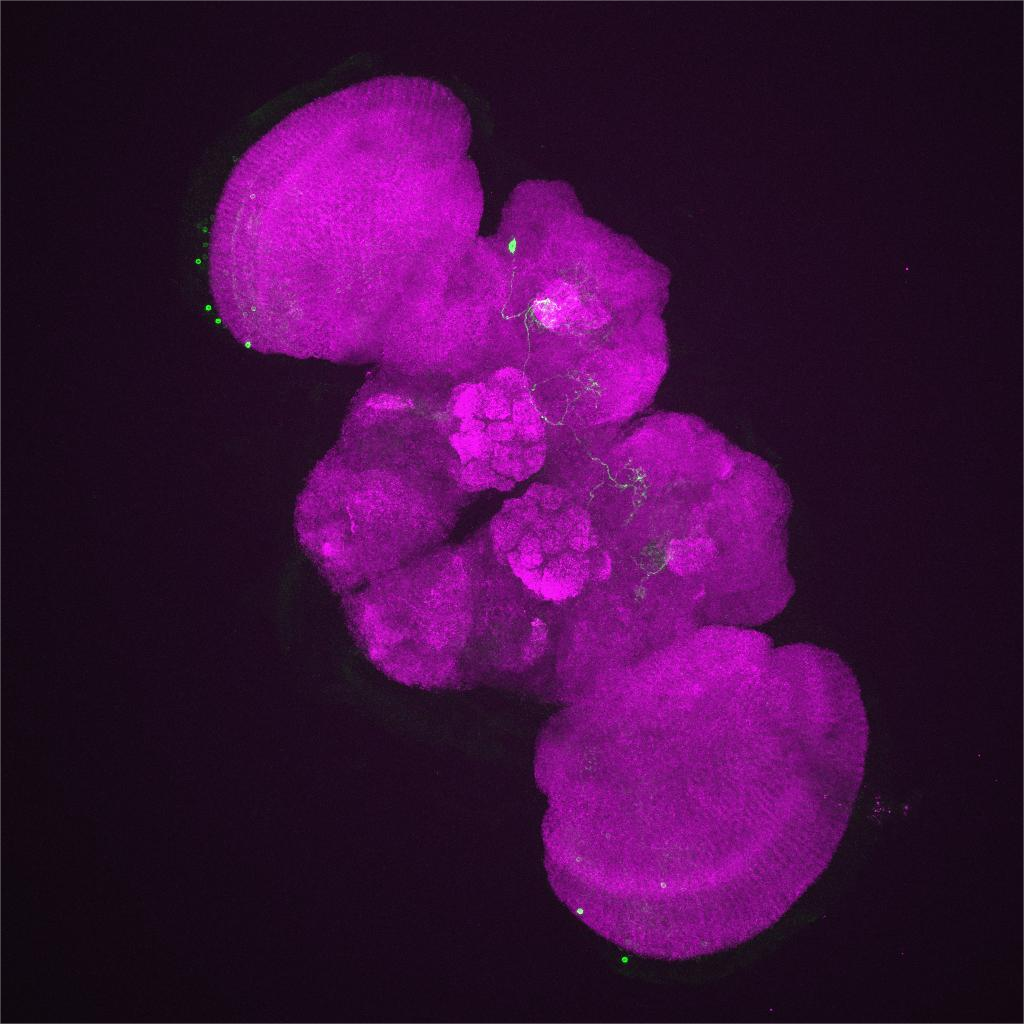 |
| R64H01 | *GAL4*^R64H01^*/*  *UAS-sNPF-R-RNAi* | **** | <0.0001 | 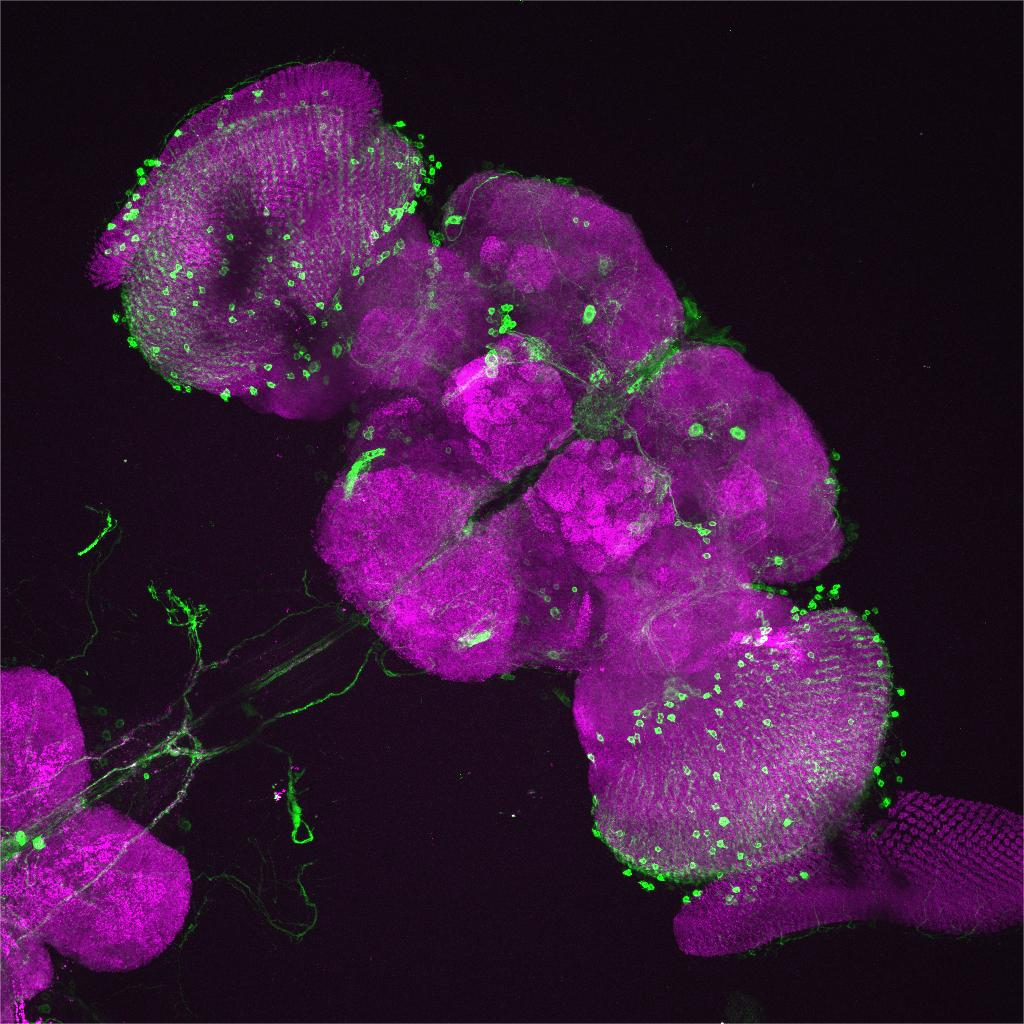 |
